# Spatiotemporal organization of BamA governs the pattern of outer membrane protein distribution in growing *Escherichia coli* cells

**DOI:** 10.1101/2021.01.27.428258

**Authors:** Gideon Mamou, Patrick G. Inns, Dawei Sun, Renata Kaminska, Nicholas G. Housden, Ruth Cohen-Khait, Helen Miller, Kelly M. Storek, Steven T. Rutherford, Jian Payandeh, Colin Kleanthous

## Abstract

The outer membrane (OM) of Gram-negative bacteria is a robust protective barrier that excludes major classes of antibiotics. The assembly, integrity and functioning of the OM is dependent on β-barrel outer membrane proteins (OMPs), the insertion of which is catalyzed by BamA, the core component of the β-barrel assembly machine (BAM) complex. Little is known about BamA in the context of its native OM environment. Here, using high-affinity fluorescently-labelled antibodies in combination with diffraction-limited and super-resolution fluorescence microscopy, we uncover the spatial and temporal organization of BamA in live *Escherichia coli* K-12 cells. BamA is clustered into ~150 nm diameter islands that contain an average of 10-11 BamA molecules, in addition to other OMPs, and are distributed throughout the OM and which migrate to the poles during growth. In stationary phase cells, BamA is largely confined to the poles. Emergence from stationary phase coincides with new BamA-containing islands appearing on the longitudinal axis of cells, suggesting they are not seeded by pre-existing BamAs but initiate spontaneously. Consistent with this interpretation, BamA-catalyzed OMP biogenesis is biased towards non-polar regions. Cells ensure the capacity for OMP biogenesis is uniformly distributed during exponential growth, even if the growth rate changes, by maintaining an invariant density of BamA-containing OMP islands (~9 islands/μm^2^) that only diminishes as cells enter stationary phase, the latter change governing what OMPs predominate as cells become quiescent. We conclude that OMP distribution in *E. coli* is driven by the spatiotemporal organisation of BamA which varies with the different phases of growth.

**Significance:** The integrity and functioning of the outer membrane (OM) of Gram-negative bacteria depends on the β-barrel assembly machinery (BAM). Although the structure and the mechanism of the complex have been widely explored, little information exists about the organization of the BAM complex and how it dictates protein distribution in the OM. Here, we utilized highly specific monoclonal antibodies to study the spatiotemporal organization of BamA, the key component of this complex. We reveal that BAM organization is dynamic and tightly linked to the cell’s growth phase. We further discover that the rate of BAM facilitated OMP biogenesis is significantly reduced near the poles. In turn, these features govern the biogenesis patterns and the distribution of OMPs on the cell surface.

## Introduction

The outer membrane (OM) of Gram-negative bacteria is the outermost layer of the envelope which serves as the interface between the cell and the surrounding environment (1). The OM is an asymmetric bilayer composed of an inner leaflet of phospholipids and an outer leaflet of lipopolysaccharides (LPS) (2, 3). This architecture protects the cell from detergents and antimicrobial agents (4–6) and its rigidity plays an important role in maintaining cell morphology (7). Embedded within the OM is an assortment of β-barrel outer membrane proteins (OMPs) which carry out essential roles such as building and maintaining the membrane, adhering to host epithelia during bacterial pathogenesis, bringing nutrients into the cell - both passively and actively - and forming numerous stabilizing attachments to the underlying cell wall (2, 8–11). OMPs are inserted into the membrane by the β-barrel Assembly Machine (BAM) (12, 13), which is essential in Gram-negative bacteria. An equivalent machine is responsible for the insertion of OMPs into plastid and mitochondrial OMs (14). Unsurprisingly, given its central role in the cell envelope, the BAM complex is an important antibiotic target (15–18).

The BAM machinery is a ~200-kD complex consisting of four periplasmic lipoproteins (BamB-E) and BamA, an OMP (19–21). In addition to its 16-stranded β-barrel, BamA also has a periplasmic extension of 5 POTRA domains, which, together with the other BAM components, catalyze insertion of OMPs into the membrane (12, 13). Key mechanistic facets of BamA known to be important for its function are its lateral gate, the non-covalent junction between its first and last β-strands, and loop 6 which caps the central cavity of the barrel (22–24). The periplasmic lipoproteins seem to play important roles in regulating the flow of substrates to BamA and its conformational transitions (25–29). Although the structure and function of the BAM complex has been studied extensively (19–21), the mechanism of BAM-mediated OMP insertion is still being elucidated (30, 31). A ‘membrane thinning’ model argues that the asymmetric thickness of the BamA β-barrel thins the OM close to the lateral gate thereby facilitating the spontaneous insertion of OMPs in the OM. The budding model suggests that BamA plays an active role in OMP folding by templating the folding process itself on its lateral gate β-strands. Evidence in support of both models has been presented (32–36).

Compared to the numerous *in vitro* investigations of BAM-mediated OMP folding, few studies have explored the spatial organization of OMPs and their biogenesis sites in live bacteria (37–39). Recent work has shown that OMPs are clustered into islands that are distributed throughout the OM (40–43). Supramolecular organization is the means by which Gram-negative bacteria passively turnover the protein composition of the OM through multiple divisions since old OMPs are predominantly localized to cell poles while new OMPs are mostly located at midcell (44). There has been no similar study of BamA localization and consequently we have little understanding of how OMP organization and biogenesis are linked. Here, we characterize the spatiotemporal organization of BamA in live cells using high-specificity monoclonal antibodies. We show that BamA forms islands containing multiple BamA molecules that are distributed across the OM and that migrate towards the poles as cells grow. The spatial organization and density of BamA-containing islands is tightly regulated during exponential growth, but changes as cells enter stationary phase. By comparing the distribution of OMP biogenesis with BamA organization, we show that changes to BamA localization are linked directly to OMP insertion, which in turn dictates how the OM is populated with OMPs during growth.

## Results

### Labelling BamA in Live Cells

Exploring how BamA is organized is essential for understanding how the OM is populated with OMPs. A previous study, where BAM proteins were imaged at a sub-diffraction resolution, showed that BamA forms clusters on the surface of the OM in *E. coli* (43). However, this study used fixed cells that precluded the simultaneous visualization of BamA and newly-inserted OMPs in live bacteria. Existing structural information shows that most of the BAM complex resides in the periplasm while the most surface-exposed protein regions are the extracellular loops of the BamA β-barrel (19–21). In order to label these residues, we used a high-affinity monoclonal antibody (MAB2) that binds specifically to loop 6 of BamA without impacting function (15). To maximize our ability to label the surface-exposed regions of BamA molecules, fluorescently-tagged MAB2 fragment antigen-binding (Fab) domains were used. We first confirmed that cell growth and viability were unaffected by this Fab (Figure S1A). We then used the label to visualize BamA in live *E. coli* cells. Both epifluorescence and 3D-SIM images showed that BamA forms clusters on the surface of the membrane, resembling previously described OMP islands (Figure 1A, Movie S1) (44). Labelling efficiency was enhanced when we used a deep-rough (*waaD*) strain which has a minimal LPS structure and thereby allows greater accessibility of surface-exposed epitopes (Figure S1B). Previous studies have shown that this mutant is viable and produces similar amounts of BamA (45, 46). This mutant was used for many of the experiments reported here. To validate the specificity of our labelling, we used a strain where the sequence of the BamA β-barrel was replaced with that from *Klebsiella pneumoniae* (15). In this case, no BamA labelling was observed although cell viability and the formation of OMP islands (imaged through labelling the vitamin B_12_ transporter, BtuB) on the cell surface were unaffected (Figure S1C).

**Fig 1.**
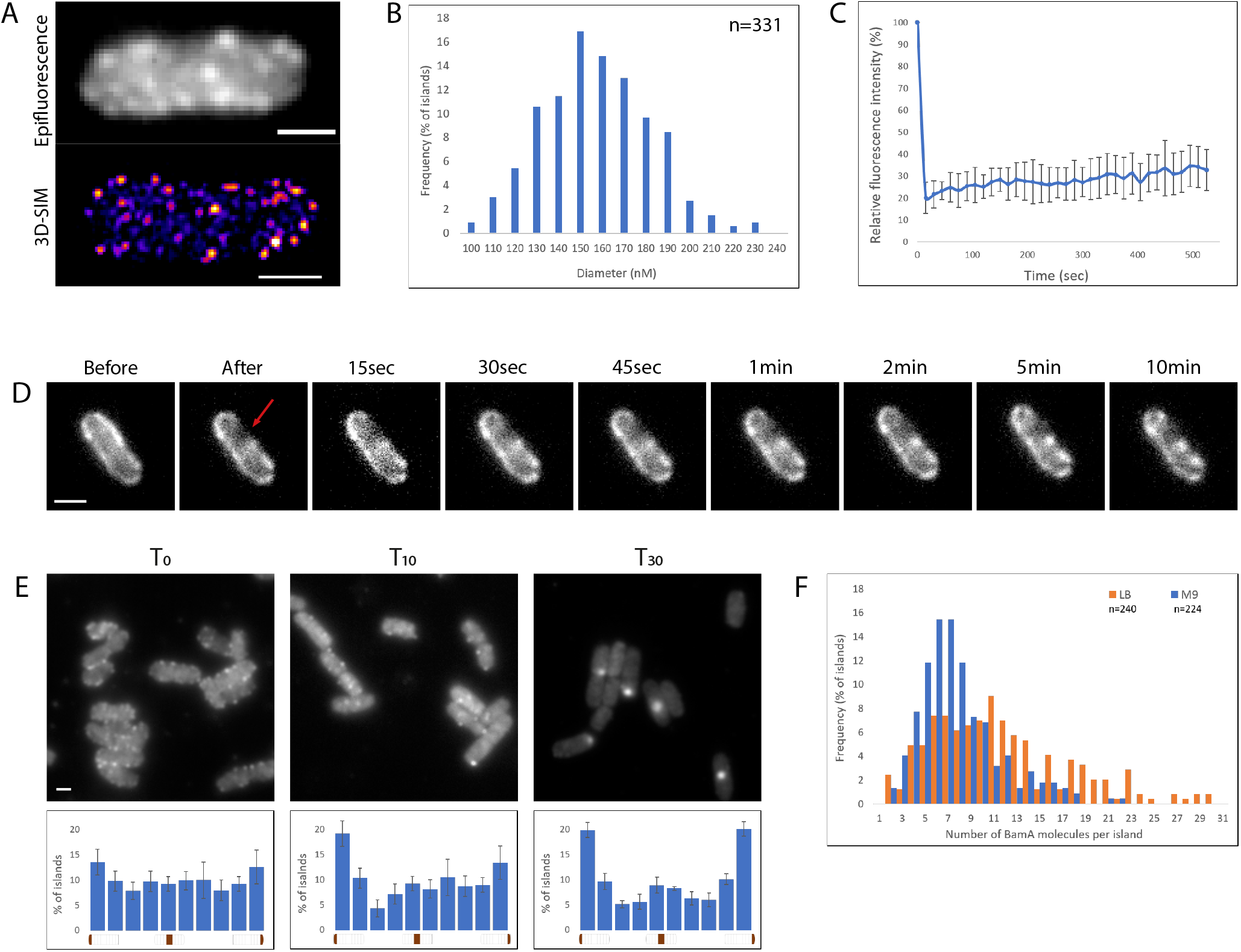
**(A)** Images of live, single *waaD E. coli* cells labelled with αBamA^AF488^ Fabs. **(B)** Size distribution of BAM islands based on 2D-SIM images. The average diameter of an island was 153±25nm. **(C)** FRAP plot of BamA in live cells demonstrating its low mobility in the membrane. Calculations are based on 4 biological repeats. **(D)** Representative time-lapse images of a BamA FRAP experiment. Red arrow indicates the bleached region. **(E)***E. coli* cells were labelled with αBamA^AF488^ Fabs at T_0_, and resuspended in fresh LB media for the indicated amount of time. Shown are representative images of Bam-containing islands (*Top panels*) and distribution plots from biological triplicates (*Bottom panels*) at the indicated time points. For T_30_, at least one doubling time was confirmed by OD_600_ measurement. **(F)** The number of BamA molecules within an island based on Single Step Photobleaching measurements. Shown is the distribution of BamA copy number per island in LB vs. M9 media. The average number of molecules was 11.1 (LB) and 6.5 (M9). Scale bars, 1 μm.

### Imaging the Spatial Organization of BamA in Growing Cells

After establishing a protocol for rapid labelling of BamA, we explored its surface architecture and mobility in live cells. The addition of antibiotics that block transcription or translation led to a decrease in the fluorescence intensity of BamA-containing islands, consistent with reduced levels of BamA in the OM (Figure S1D). Cells growing in different media types displayed clustered distribution of BamA, but the amounts of protein varied, as deduced by both fluorescence imaging and Western blotting of OM extracts (Figure S1E-G). Measuring the dimensions of BamA islands by SIM microscopy showed that their size varies; the average diameter was ~150nm (Figure 1B). These dimensions were similar in different bacterial strains, growth media and when cells were fixed (Figure S1H). Similar BamA island dimensions were reported using fixed cells (43). Next, we sampled the mobility of BamA in the OM using Fluorescence Recovery After Photobleaching (FRAP) experiments, which demonstrated that BamA is essentially immobile in the OM; less than 35% of the signal recovered over a 10 min time-course (Figure 1C-D, Movie S2). The lack of recovery, which was independent of the region being bleached (Movie S3), is consistent with previous FRAP studies on other labelled OMPs (BtuB and Cir) that are known to reside within OMP islands (44). We went on to track the movement of BamA-containing islands as cells grew and divided. Cells that were imaged immediately after labelling displayed a uniform distribution of islands across the long axis of the cell (Figure 1E). When these cells were resuspended in fresh media and imaged after 10 or 30 minutes, the distribution changed significantly, with most BamA-containing islands localizing to polar regions (Figure 1E, Figure S2A). To confirm the change in distribution was attributed to cell growth, we carried out a similar experiment in the presence of rifampicin. In contrast to fast dividing cells, the distribution of BamA in the treated cells remained uniform (Figure S2B). Together, these results indicate that BamA migrates towards cell poles in a growth-dependent manner. Similar observations have been made for BamC (44). We conclude therefore that the entire BAM assembly migrates to the poles as cells grow.

One of the advantages of our imaging strategy is that BamA molecules are labelled by a single Fab. Thus, we were able to use Single Step Photobleaching (SSP) microscopy to estimate the number of BamA complexes within a cluster (Figure S2C-D, Movie S4). Data from multiple experiments showed that for cells grown in LB, the average number of BamA molecules per island was 10.5 (Figure 1F). This number varied within and among cells as some islands contained only 3-4 copies while others contained more than 20. When similar measurements were applied to cells growing in M9 media, where the growth rate was lower, the number of BamA molecules per island ranged between 3 and 15 with an average of 6.5 molecules (Figure 1F). Since the diameter of the observed islands was typically ~150 nm (Figure 1B), it is likely that they contain both BamA and an assortment of other OMPs. Using 3D-SIM images, we also determined that the density of the islands was approximately 8-10/μm^2^ (Figure 2A) and therefore we estimated the copy number of BamA on the surface as ~700-800 proteins per cell when grown in LB. These values are in accord with previous proteomics studies (47, 48).

**Fig 2.**
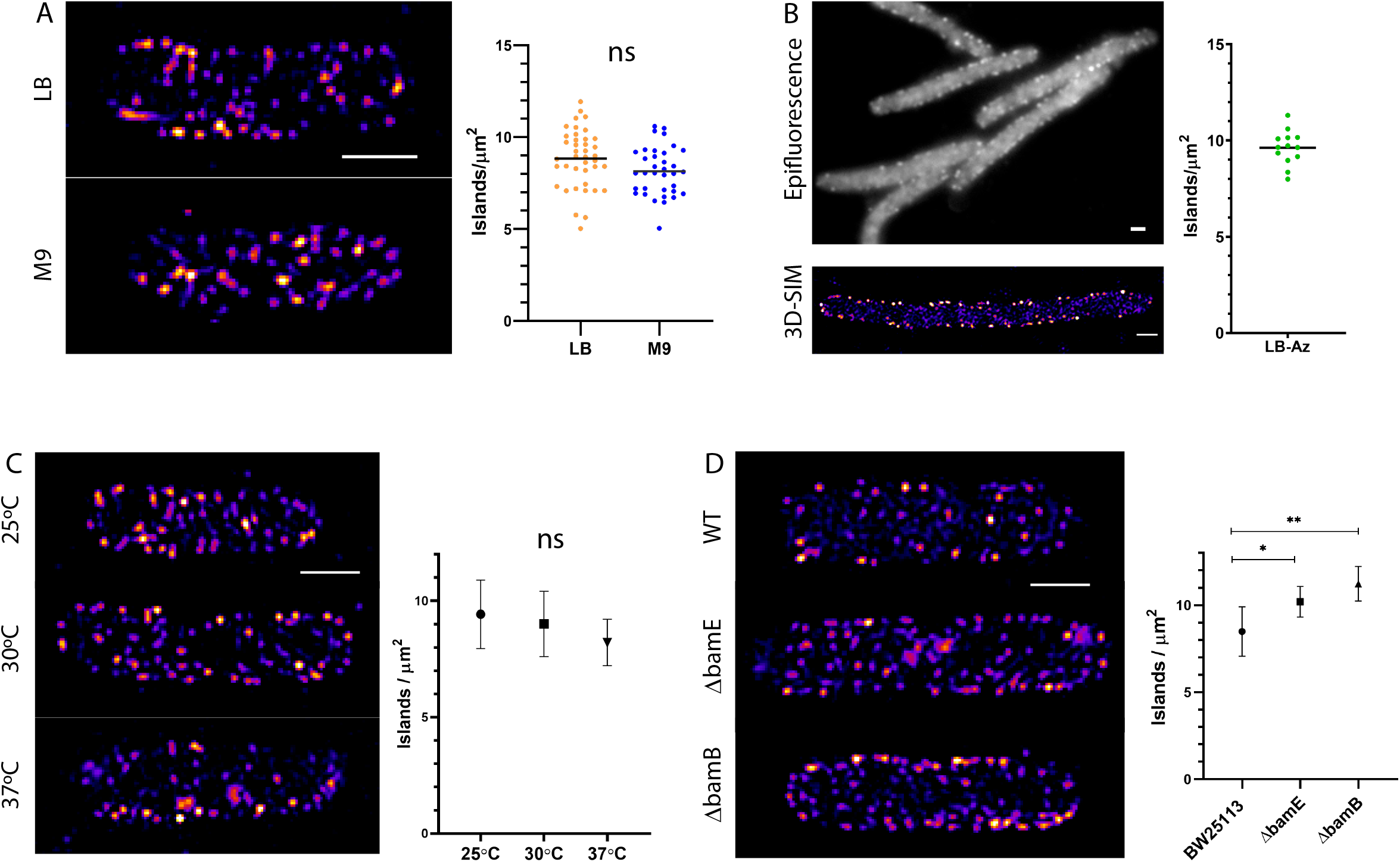
All panels show representative 3D-SIM projections of BamA and corresponding measurements of the density of BamA-containing islands on the surface of *E. coli*. **(A)** A comparison of cells grown in LB vs. M9 media. Samples from both media types were taken at a similar OD_600_ (mid-log phase). **(B)** Shown are cells grown in LB media treated with 1 μg/ml aztreonam. The antibiotic was added to exponentially growing cells 30 minutes before sampling, which had little effect on the OD_600_ compared to non-treated cells. **(C)** Comparison of cells grown at three different temperatures. Samples were taken at a similar growth stage (See figure S3C). **(D)** Comparison of wild-type *E. coli* K-12 (BW25113) and two BAM lipoprotein deletion strains. Cells were grown in LB and samples taken at a similar OD_600_. Scale bars, 1 μm.

### The Density of BamA-Containing Islands Remains Constant During Cell Growth

As bacterial cells grow, new BAM complexes need to be incorporated into the expanded OM to maintain the cell’s requirement for new OMPs. To determine whether the density of BamA-containing islands is affected by alterations in growth rate, we compared the density between cells grown in LB and M9 media. Surprisingly, we found that the density is similar in both media types despite the fact that lower amounts of BamA were found in M9-grown cells (Figure 2A, Figure S1E-G). We then investigated whether the density is maintained when cell division is perturbed, following treatment with the PBP3 inhibitor aztreonam which causes cells to filament. Although cells became much longer, the density of the BamA-containing islands on the cell surface remained largely unchanged (Figure 2B), suggesting the insertion of new BamA-containing islands in the OM is coordinated with cell elongation. Epifluorescence microscopy and Western blot analysis showed that aztreonam and cephalexin treatment did not significantly affect the overall levels of BamA in the OM (Figure S3A-B). Next, we grew cells at various temperatures to test whether this would influence island density. Once again, the density of BamA-containing islands did not vary dramatically despite the change in environmental conditions (Figure 2C); even as cells grew faster at higher temperature (Figure S3C) there was no significant increase in the amount of BamA detected either by epifluorescence microscopy or Western blotting (Figure S3D-E). Our results demonstrate that the density of BamA-containing islands in the OM of exponentially growing *E. coli* is maintained while the copy number of BamA molecules per island can vary.

OMP islands are thought to be stabilized by promiscuous protein-protein interactions (41, 49). It has recently been suggested that specific associations of BamB from two neighbouring BAM complexes might explain the propensity for the biogenesis machinery to cluster in the OM (43), and consequently drive formation of OMP islands. We therefore tested whether the size or density of BamA-containing islands were altered in *bamB* or *bamE* deletion strains. We found that in both strains, BamA formed clusters across the cell surface (Figure 2D, Movie S5), indeed the density of the islands was slightly higher compared to the parent BW25113 strain. We also prepared OM extracts from these strains to measure the total amount of BamA in the OM. Unlike the microscopy data, the OM of the *bamB* strain contained 35±22% less BamA than the wild-type strain although the amount varied between biological replicates (Figure S3F). Since deletion of *bamB* has been shown to affect both the efficiency of the BAM complex and proteinaceous content of the OM (27, 50), we investigated whether *bamB* and *bamE* deletions affected the insertion of the OMP BtuB. While high expression level and formation of BtuB clusters were clearly observed in the OM of the wild-type strain, the number and fluorescence intensity of the islands was significantly reduced in both mutant backgrounds (Figure S3 G-H). The effect was more severe in the *bamB* deletion strain where very little BtuB was detected and mostly localized near the poles. Taken together, our data suggest that deletion of the BamB lipoprotein affects the capability of the complex to insert certain OMPs into the OM but has little influence on BAM clustering.

### BamA Organization Comprehensively Changes at Stationary Phase

As bacterial cell density increases, the population begins to enter stationary phase in which profound physiological changes occur, including changes in the OM (51–53). We examined whether BamA organization changes in stationary phase cells. Strikingly, we found that cells from an overnight culture displayed very different BamA organization (Figure 3A). Most obviously, far fewer islands were observed on the cell surface in both wild-type and *waaD* strains (Figure 3A, Figure S4A-B). Similar results were observed when a different α-BamA Fab, which targets loop 4, was used (Figure S4C) (15). In addition to the decrease in the number of islands per cell, SSP experiments showed that the number of BamA molecules per island was reduced by almost 60% compared to exponentially grown cells (Figure 3B). Importantly, islands in stationary phase cells were no longer uniformly distributed across the entirety of the cell surface (Figure 3A). Integrating the localization of islands from a population of stationary phase cells clearly indicated that they were mostly localized in polar regions (Figure 3C). To confirm the apparent polar localization of BamA in stationary phase was not an artefact of epitope accessibility, for example due to changes in cell surface topology (52), cells were co-labelled with both αBamA^AF488^ and ColN^−mCherry^ (54) which binds with high affinity to OmpF. Although OmpF distribution was slightly polar biased, it was labelled across the cell and the correlation between the fluorescent intensities of the two labels was poor (Figure S4D-E), suggesting that labelling accessibility was not the reason for the non-uniform distribution of BamA. Interestingly, when the spatial distribution of other OMPs was investigated, they displayed different degrees of polarity (Figure S4F). BtuB and FepA are 22-strand β-barrel, TonB-dependent transporters (TBDTs) that are substrates of BamA (55, 56). While FepA is distributed across the surface in stationary phase cells, BtuB is largely polar-localized. We surmise that this difference in surface distribution, and indeed in fluorescence labelling intensity, likely reflects known differences in the expression of these OMPs as cells enter stationary phase, a point we return to below.

**Fig 3.**
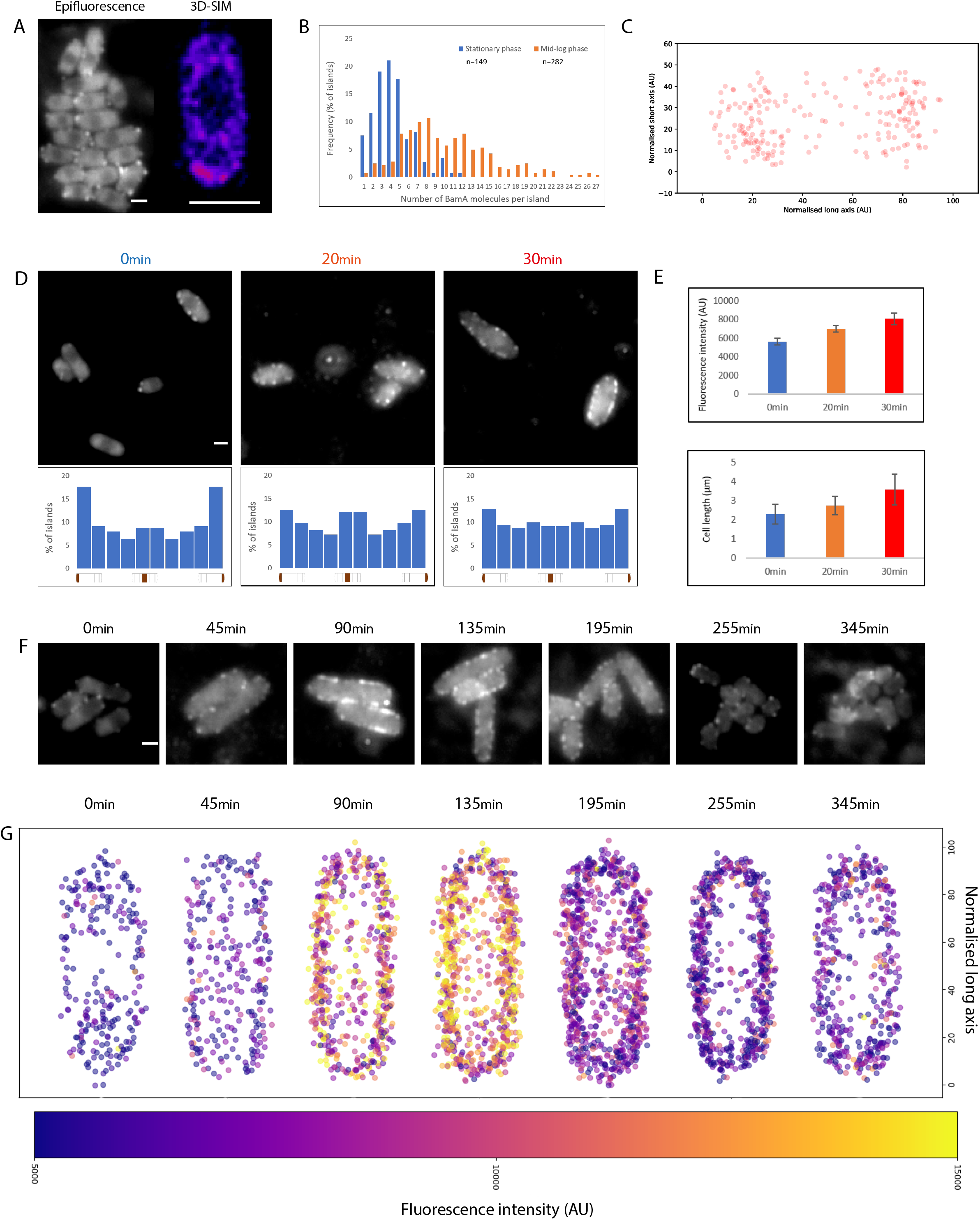
**(A)** Images of BamA labelling from an overnight culture of *waaD E. coli* cells grown in LB. **(B)** The number of BamA molecules within an island is based on Single Step Photobleaching measurements. Shown is the BamA copy number per island in stationary phase vs mid-log cells. The average number of molecules was 10.1 (LB) and 4.3 (M9), respectively. **(C)** Integrated localization map of BamA-containing islands from multiple cells demonstrating its biased polar distribution (n=221 islands). **(D)** Recovery of BamA organization after resuspension in fresh media. Cells from an overnight culture were immediately labelled (0 min) or resuspended in fresh LB for the indicated time period. Shown are representative images of BamA labelling (*top panels*) and distribution plots (*bottom panels*) (n_t0_=181, n_t10_=262, n_t30_=165 islands). **(E)** The average fluorescence intensity of BamA-containing islands and the length of cells during the recovery shown in panel D. **(F)** Time course images of BamA labelling at different growth stages. Cells were fixed immediately after sampling. Shown are representative images from the indicated time points. **(G)** Integrated localization map of BamA-containing islands from multiple cells at different growth stages. Islands are color coded according to their fluorescence intensity. Scale bars, 1 μm.

Since OMP biogenesis is essential for cell elongation and growth, bacteria need a substantial number of BAM complexes when cells emerge from stationary phase. In accord with this assertion, the amount of BamA on the surface quickly increased and its organization changed when stationary phase cells were resuspended in fresh media (Figure 3D, Figure S4G). BamA was synthesized *de-novo* as no increase in amount or organization of BamA occurred when chloramphenicol was added to the fresh media (Figure S4H). This change was evident even before the cells started to divide or elongate (Figure 3E), which supports the idea that BamA is among the first OMPs to appear in the OM after resumption from growth arrest. The observed restoration of uniform distribution implies that most new BamA molecules are inserted near midcell despite the deficiency in pre-existing BAM complexes in these regions. To examine this effect further, we compared the organization of BamA during multiple growth stages. The microscopy images revealed a trend where BamA levels constantly increase after the lag-phase until they peak during exponential growth (Figure 3F, Figure S5A-C). BamA organization remained steady until the culture became dense and the growth rate started to decrease. After this point, cells returned to the “stationary configuration” within 2-3 hrs. Integration of both the fluorescence intensity and localization of BamA-containing islands from a population of cells further supported observations from individual cells (Figure 3G). In addition, labelling LptD, the essential OMP that deposits LPS in the OM (10), showed that it too forms clusters (Movie S6) and its arrangement in the OM is also growth stage dependent (Figure S5D).

Finally, we examined whether the density of BamA-containing islands is the same or different to that in exponentially growing cells. In line with the results above, stationary phase cells displayed a 67±7% decrease in island density compared to mid-log phase cells. (Figure S5E&F). These results lead us to conclude that the spatiotemporal organization of the BAM complex is intimately linked to the growth phase of the cell.

### BamA Activity Depends on its Localization on the Cell Surface

Insertion of new β-barrel OMPs from the periplasm to the OM is carried out exclusively by the BAM complex (12, 13), with recent studies suggesting that a direct link exists between the Sec secretion machinery in the inner membrane and BAM in the OM (57–59). Several studies have suggested that OMP biogenesis takes place across the entire membrane (37, 39, 43) while others argue that certain areas are more permissive than others (38, 39, 44). However, none of these studies have monitored both OMP biogenesis and BamA localization simultaneously. To visualize new OMPs appearing on the surface we used strains expressing FepA or BtuB from an inducible plasmid and labelled them with fluorescently-labelled colicins, which bind the OMPs with high affinity and specificity (Figure S6A&D). Using this assay, we could detect FepA appearance on the surface after just 3 minutes of arabinose induction (Figure 4A). We confirmed that FepA was *de-novo* expressed as chloramphenicol treatment negated its insertion into the membrane even after a longer induction (Figure S6B). After 5 minutes induction, the spatial distribution of FepA insertion was clear and we could see that FepA is mostly present at midcell (Figure 4B, Movie S7). Quantification of the fluorescence intensity from multiple cells further supported the observation that polar regions are frequently devoid of new FepA while midcell is the most populated (Figure 4C). Since OMP diffusion is very slow and the cells were fixed immediately after sampling, it is reasonable to assume that the observed localization of FepA represents the original site of insertion. Midcell-biased biogenesis was also observed when cells were grown in M9 media and when the induction time was increased to 15 minutes (Figure S6C). Similar biogenesis patterns were observed when BtuB was expressed briefly and labelled using fluorescently labelled ColE9 (Figure S6D-F, Movie S8). Intriguingly, we noticed that in some cases, FepA appeared only in a single pole. We therefore inspected whether new FepA appears at the poles after different induction periods (Figure 4D). At 5 minutes post induction, most cells did not have FepA at the poles or displayed unipolar labelling. When the induction time was increased, the bipolar pattern became more common. The fact that unipolar labelled cells tended to be short cells implies that this pattern results from midcell-biased biogenesis in cells which then divided during induction. The increase in bipolar labelling frequency after 20 and 40 minutes might result from either cell division or polar migration of the OMPs. Next, we tested whether observed changes in BamA levels had an impact on OMP biogenesis. Although our measurements showed that in LB media BamA-containing islands have ~50% more BamA than in M9 there was only a slight difference in biogenesis levels detected (Figure S6G). Similar comparisons between wild-type and deep rough LPS mutant strain, in which its activity is moderately compromised (46), also did not show a major difference (Figure S6H). We suggest these observations point to the possibility that OMP insertion into the OM is not a bottle neck and that the multiple BAM assemblies within islands may be somewhat underused.

**Fig 4.**
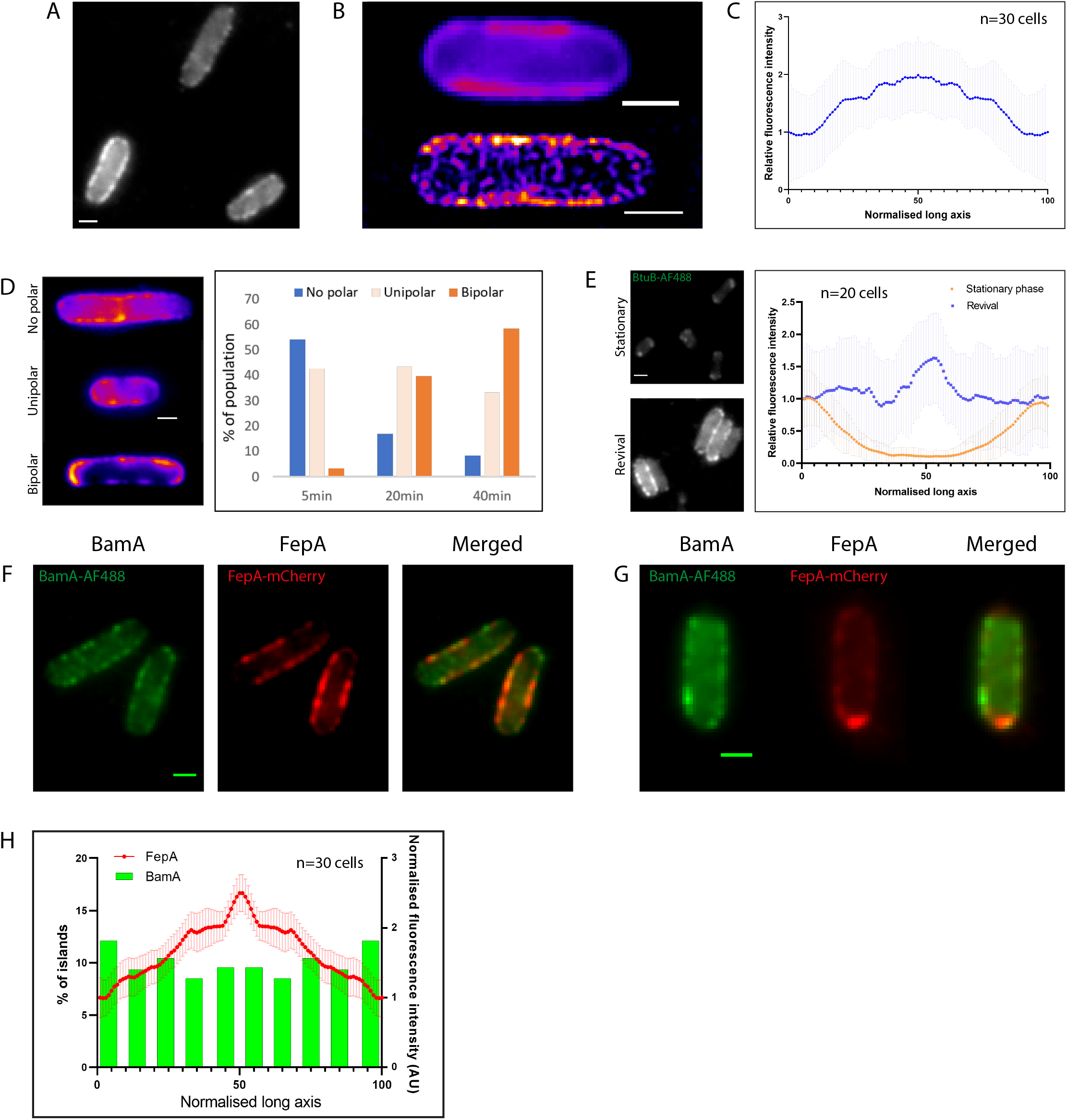
**(A)** GM07 *E. coli* cells labelled with ColB-GFP three minutes after induction of FepA, expressed from a plasmid. Cells were fixed immediately after sampling. **(B)** Representative images of midcell-biased FepA biogenesis 5 minutes after induction. Shown are epifluorescence (*top*) and 3D-SIM projection heatmaps (*bottom*). **(C)** A kymograph of normalized fluorescence intensity profiles of FepA based on the integrated analysis of multiple cells after 5 minutes induction. **(D)** Representative heatmaps of the three main FepA distribution patterns observed (*left*) and the abundance of each pattern after different periods of induction (*right*). **(E)** Representative images (*left*) and a kymograph of normalized fluorescence intensity profiles of BtuB (*right*) at stationary phase vs revival in fresh media. **(F)** Co-labelling of BamA and FepA after a short induction of FepA biogenesis. Shown are epifluorescence images highlighting the discrepancy between the localization of BamA and the lack of corresponding biogenesis at the poles. **(G)** Co-labelling of BamA and FepA in cells where new FepA is located at one pole. Shown are epifluorescence images demonstrating that this pattern is not the result of BamA localized at the poles. **(H)** Comparative analysis of BamA and FepA distribution from multiple cells after 7 minutes induction (n=20 cells, 235 islands). Shown are the distribution of BamA-containing islands (*green columns*) and the normalized fluorescence intensity profile of FepA (*red line*). The same cells were used for both types of analysis. Scale bars, 1 μm.

Our OMP induction experiments clearly show that OMP biogenesis is midcell-biased. We next investigated whether this pattern of biogenesis is also the case for natively expressed OMPs. We compared the distribution of BtuB and FepA in stationary phase and after a short resuspension in fresh media (revival experiments). The distribution of BtuB, which is known to be expressed during exponential growth (60), was transformed from being completely polar to being midcell-biased well before cells started to divide (Figure 4E, Movie S9). The overall amount of BtuB in the OM was also significantly increased (Figure 4E, Figure S7B) indicating that BtuB biogenesis is the reason for this change. In contrast, the distribution of FepA, which is mostly expressed during later growth stages (60, 61), became more polar after cell revival (Figure S7A, Movie S10). Co-labelling of both OMPs before and after revival revealed a striking spatial compartmentalization and re-organization. In stationary phase, FepA was dominant and occupied most of the midcell region, but after revival, FepA was pushed to the poles while the newly inserted BtuB inhabited midcell (Figure S7C). These results confirm that OMP biogenesis is centred around midcell and that polar migration is driven by growth.

Previous studies have suggested that OMP biogenesis is unequally distributed along the *E. coli* cell surface. However, these studies had not simultaneously monitored both BamA localization and OMP biogenesis in the same growing cell. Exploiting the approaches, we have developed, we addressed this question by co-labelling cells for BamA and FepA, the latter following a brief period of induction. While BamA-containing islands were distributed throughout the cell (as described above), FepA biogenesis was largely restricted to midcell. Much less FepA was detected near the poles despite the presence of BamA (Figure 4F, Figure S7D). We note that in short cells, which must have recently divided, BamA was uniformly distributed while OMP biogenesis showed a distinct unipolar distribution (Figure 4G). Quantitative analysis at the population level revealed that despite the lower biogenesis rates at the poles, the abundance of BamA was similar to other cellular regions (Figure 4H). These results show definitively that although BamA-containing islands are found throughout the OM those islands residing near the poles have much lower OMP biogenesis activity than those at midcell.

## Discussion

Our data demonstrate that the organisation of BamA in elongating cells affects OMP patterning in the OM (Figure 5). In exponentially growing cells, BamA-containing islands are distributed uniformly across the OM. These islands contain an average of 10-11 BamA molecules and are large enough to accommodate many other OMPs, which are likely to be BamA substrates that have not diffused far from their progenitor BamA (41). As cells grow, new OMP islands are inserted in non-polar regions, pushing old islands towards the poles, as previously proposed (41, 44). Thereafter, the insertion of new BamA-containing islands is coordinated with cell elongation such that the density of islands remains constant. When cells divide, the non-uniform distribution of OMP biogenesis creates an OMP arrangement in which the old pole contains old OMPs and the new pole new OMPs. After several division cycles, cell density increases and entry into stationary phase triggers a major change in both BamA organization and the OMP content of islands. As a consequence, BamA becomes predominantly polar-localized. At this stage, polar islands contain a mixture of old, mid-log and stationary phase expressed OMPs while the more centrally localized islands are populated with the last OMPs to be expressed during the latter growth stages.

**Fig 5.**
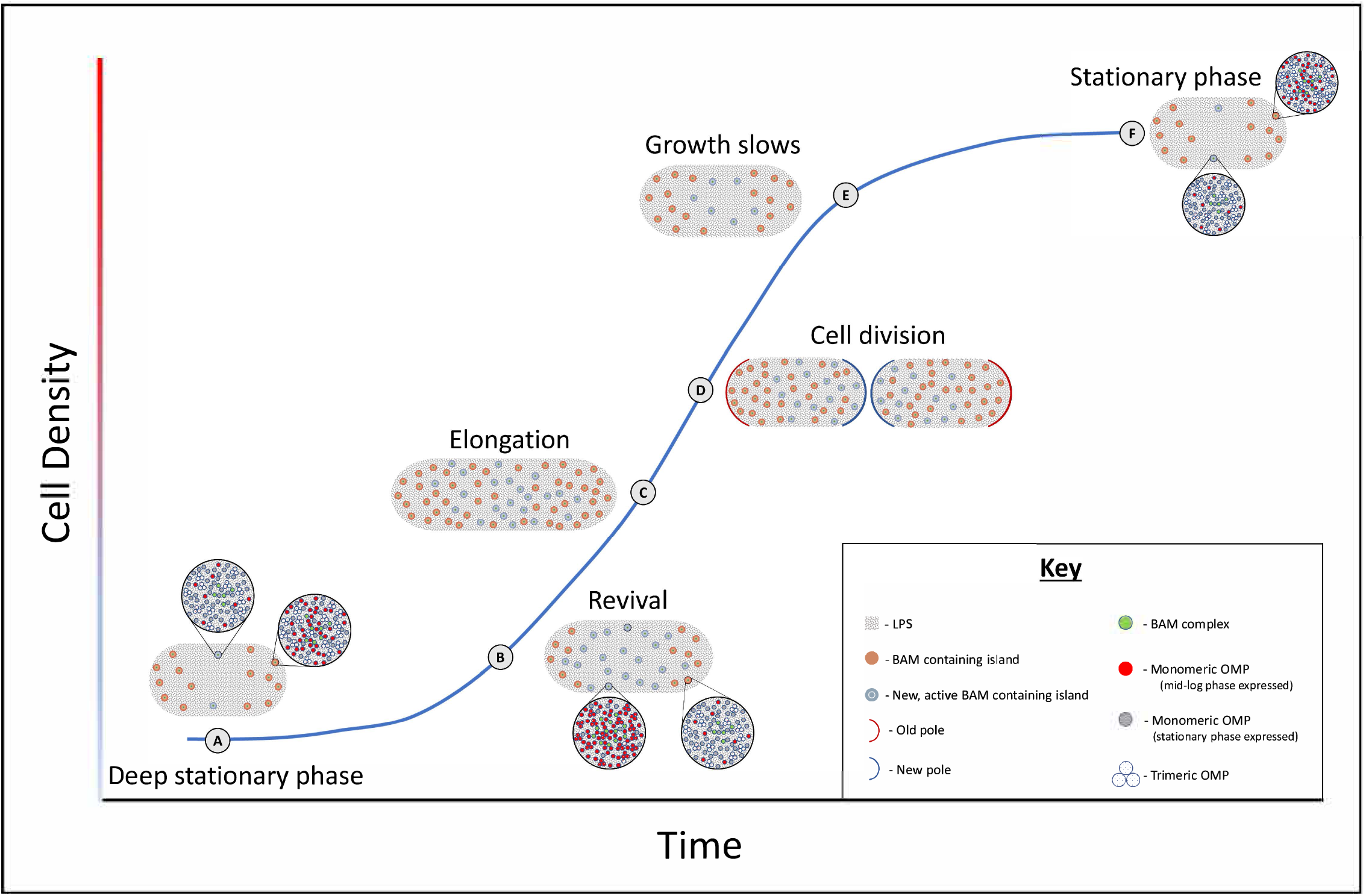
Model depicting how the spatiotemporal changes in BamA during growth govern OMP organization in the OM. (**A**) in deep stationary phase cells, BamA is mostly located at the poles and the content of the islands varies between those at midcell and the poles. (**B**) Shortly after the emergence from stationary phase, many new BamA containing islands appear on the long axis of the cell, pushing old OMPs towards the poles. (**C**) Mid-log cells maintain a constant density of BamA-containing islands and exhibit midcell-biased OMP biogenesis. (**D**) Following cell division, new OMPs are located at new poles while older OMPs localize to old poles. (**E**) As cell density increases, growth slows, leading to a decrease in BamA expression and a shift in OMP expression. (**F**) The last few cell divisions re-establish the stationary phase configuration and this OMP arrangement is maintained until cell growth resumes.

BamA organization changes profoundly on entry into stationary phase. The changes include a decrease in the density of BamA-containing islands, a drop in BamA copy number per island and redistribution across the OM so that few BamA-containing islands are at midcell and those that remain are polar localized. Comparing the spatiotemporal organization of BamA upon entry into stationary phase and after 24 hours indicates that these changes occur shortly after the growth rate begins to slow and are almost completed by the beginning of stationary phase. Previous studies have shown that the transition to stationary phase is the period where extensive switches in OMP expression take place as well as many regulatory changes, such as increased expression of chaperones and the *rpoE* regulon. (61, 62). BamA expression decreases as cells stop growing (60, 61, 63), with the few remaining divisions diluting the existing BAM complexes and pushing them towards the poles. OMPs such as BtuB, Cir and LamB are known to migrate to the poles under the force of growth (40, 44). The present work shows that the OMP biogenesis machine is on the same default pathway to the poles of an elongating cell. Moreover, LptD, which deposits LPS into the OM, is on the same default pathway and undergoes similar organizational changes on entry into stationary phase.

In conjunction with other studies, our data suggest that BamA may have two routes into the OM of *E. coli*; one that depends on pre-existing BamA within islands and one that may be BamA-independent. The BamA-dependent insertion of other BamA proteins is well-established and is supported by recent cryo-EM structures of BamA caught-in-the-act of folding another BamA molecule (36). Such BamA-dependent BamA insertion explains how multiple BamA proteins are found within islands.

Two pieces of indirect evidence support the existence of a BamA-independent mode of BamA entry into the OM. First, BamA molecules within islands are immobile and so unlikely to diffuse in the membrane to seed new islands. Second, new BamA-containing islands appear early on the long axis of cells as they emerge from stationary phase despite the fact that there are few pre-existing BamAs in those regions. How could BamA be inserted into the OM without the help of pre-existing BamAs? The answer may lie in past biochemical work showing that BAM lipoproteins can aid the insertion of BamA into membranes (64, 65). If this is the case, then the seeding of new OMP islands may have their origins in the Lol pathway, which inserts lipoproteins into the OM (66). However new OMP islands are seeded, the mechanism must be closely allied to the elongation rate of cells so that a constant density of BamA-containing islands in exponentially growing cells is maintained. The mechanism must also sense the metabolic changes associated with the onset of stationary phase so that less BamA-containing islands are seeded.

It has long been debated if OMP biogenesis is uniformly distributed across the OM (37–39). By observing both OMP biogenesis and BamA localization at the same time, we observe that OMP biogenesis is strongly biased towards midcell and that this non-uniformity results from a drop in BamA activity at the poles. Our previous work showed that OMPs migrate towards the poles during cell growth (44). Here, we demonstrate that the activity of BamA depends on its OM localization. Although the molecular basis for this remains unknown, it is tempting to speculate that this could be related to substrate availability and the capability of the entire BAM complex to interact with the Sec system (59).

## Materials and methods

### Strains and plasmids used in this study

Bacterial strains and plasmids are listed in Table S1. For the construction of the pNGH206 plasmid, site-directed mutagenesis was used to introduce a solvent accessible cysteine (K469C) in the cytotoxic domain of a construct in which the N-terminal 62 amino acids of the colicin had been deleted (Δ^2-61^ ColE9). For the construction of pEE04, the *fepA* gene was amplified by PCR, cloned into a pBAD/Myc-HisB (Novagen) plasmid and transformed into either Bl21 (DE3) or BW25113 *ΔfepA* (67) *E. coli* cells.

### Antibodies and colicins used in this study

Antibodies and engineered colicins used are listed in table S2. **ColicinE9-AF488 expression and purification:** The expression and purification of this protein has been described elsewhere (41). Here, we used a modified construct with a single cysteine (Δ^2-61^ ColE9 K469C-Im9_His6_). Cys469 in the C-terminal DNase domain of these ColE9 constructs was labelled with a 3-fold Alexa Fluor 488-maleimide (Invitrogen) as previously described (41). The labelling efficiency (typically 0.8 fluorophores/protein) was estimated spectrophotometrically (V550 spectrophotometer, Jasco). **Colicin N-mCherry expression and purification:** pKBJ51 encoding Colicin N^1-185^ mCherry with a C-terminal His tag, was transformed into BL21 (DE3) (New England Biolabs). Transformed cells were grown in approximately 5L LB supplemented with 100μg/ml ampicillin. At an OD600 of approximately 0.7, induction was carried out with 1mM Isopropyl-D-thiogalactopyranoside (IPTG) for 3-4 hours at 37 °C. Cells were lysed by sonication, cell debris pelleted by centrifugation and the supernatant containing Colicin N^1-185^ mCherry extracted and filtered. Purification was carried out by conducting Ni-affinity chromatography, followed by dialysis in 25mM Tris, 150mM NaCl, pH 7.5, followed finally by size-exclusion chromatography on a 26/60 superdex 200 column (GE Healthcare). At each stage, appropriate fractions were pooled according to elution profile and SDS-PAGE analysis. Pure Colicin N^1-185^ mCherry was stored at −20 °C. **Colicin B-GFP/mCherry expression and purification:** Colicins B^1-341^ GFP/mCherry were expressed with a 6xHis tail at the C’ terminus and cloned into the multiple cloning site 2 of a pACYCDuet-1 (Novagen) plasmid where they were expressed under a T7 promotor. The plasmids were transformed into BL21(DE3) *E. coli* cells - transformed cells were grown at 37° C in LB media (pH 7.2) while shaking to OD_600_~0.6 at which point 1 mM IPTG was added and the temperature was reduced to 20° C for overnight incubation. Colicin B was purified as described in previously (70). In brief, expressing cells were re-suspended in 20 mM Tris pH 7.5, 0.5 M NaCl, 5 mM imidazole and sonicated (Sonicator 4000: 70%, 1.5 min, 3 sec on:7 sec off). The sonicated cellular extract was spun down, and the supernatant was incubated with His-binding resin (Merck 69670-5) for 10-30 min at room temperature. The Ni-resin and the bound protein were then gently (1000 x g) spun down, washed three times and re-suspended in the same buffer containing 0.5 M Imidazole which allowed protein elution. The protein was then dialyzed into PBS at 4 °C overnight. Protein concentrations were determined through absorbance at 280 nm using a sequence-based extinction coefficient.

### Expression, purification and labelling of recombinant antibodies fragments

Constructs suitable for periplasmic expression of Fab in *E. coli* and containing sequences coding for Fab fragments of MAB1, MAB2 or anti-LptD were cloned; they were transformed into 34B8 *E. coli* cells and expressed at 30°C under control of the *phoA* promoter in CRAP phosphate-limiting autoinduction medium (PMID: 12009210) supplemented with carbenicillin (50 μg/mL). After 24 hr, cells were harvested and resuspended in PBS supplemented with one complete EDTA-free Protease Inhibitor Cocktail tablet (Roche) per 50 mL of lysis buffer, lysozyme (0.125 mg/mL), and benzonase (0.01 mg/mL). The prepared suspension was microfluidized at 15,000 psi and clarified at 50,000 x g for 30 min at 4°C. The supernatant was then resolved on protein G Sepharose beads equilibrated with PBS, using 2 mL packed resin volume per original gram of cell paste. The column was washed extensively with PBS and Fabs were eluted under mildly acidic conditions (0.56 % glacial acetic acid pH 3.6). Eluted Fabs were immediately dialyzed overnight at 4°C against buffer containing 500 mM NaCl, 10% glycerol, and 100 mM Tris (pH 8.0). Fabs were further purified on an S75 16/60 gel filtration column (GE Healthcare) using PBS (pH 7.2) as the running buffer.

Fab fragments of MAB1, MAB2 and anti-LptD were labelled by using Alexa Fluor 488 Protein Labeling Kit (Thermo Fisher Scientific) following the manufacturer’s instructions. The fluorescently labelled Fabs were passed over HiPrep desalting column (GE Healthcare) to remove the excess dye. Peak fractions were collected and concentrated, and the degree of labelling determined using liquid chromatography mass spectrometry (LC–MS).

### Cell preparation for live microscopy

Overnight LB (10 g/litre tryptone, 10 g/litre NaCl, 5g/later yeast extract [pH 7.2]), supplemented M9-glucose media (0.4% (w/v) D-glucose, 2 mM MgSO_4_, 0.1 mM CaCl_2_, 1mg/ml NH_4_Cl, 0.05% (w/v) casamino acids) or Mueller Hinton II cation-adjusted broth (0.002% Tween 80) cultures were grown at 37 °C and diluted 1:100 into fresh medium with appropriate antibiotics. Stationary phase cells were collected straight from the overnight cultures. Cultures were grown at 37 °C, unless stated otherwise, to mid-log phase (OD_600_ = 0.2-0.7) and cells were centrifuged at 7000 × g for 1 min. For transcription and translation inhibition experiments, cells were treated with rifampicin (50 μg ml^−1^) or chloramphenicol (30 μg ml^−1^), respectively. For cell division inhibition experiments, cells were treated with aztreonam (1μg ml^−1^) or cephalexin (1μg ml^−1^). The antibiotics were directly added into the growing culture 40 minutes before samples were taken (unless stated otherwise). Agar pads were prepared by mixing supplemented M9-glucose medium or PBS with 1% agarose and pouring 200 μl into 1.5 × 1.6 gene frame (Thermo Scientific #AB0577) attached to the slide. For pad formation, the gene frame was sealed by a coverslip until agarose solidified. Six microliters of cells were pipetted onto the agar pad, allowed to dry and sealed with a clean coverslip. For the induction of OMPs from a plasmid, 0.4% (w/v) arabinose was added directly into the growing culture 5 minutes before samples were taken, unless stated otherwise.

For live cell labelling, an equivalent of 100μl of cells at OD_600_ = 2.5 were pelleted by centrifugation (7000 x g, 1 min), with the pellet resuspended in 100 μl of fresh supplemented M9-glucose or PBS containing 300nM fluorescently labelled Fabs or colicins. After 20 min incubation at room temperature with mixing by rotary inversion, cells were washed twice by pelleting (7000 x g, 1 min) and finally resuspended in 50 μl PBS. For imaging of fixed cells, samples were resuspended in 4% formaldehyde (in PBS) at 4°C immediately after sampling. After 20 minutes, cells were pelleted by centrifugation (7000*g*, 1 min) and labelled as described above. To improve binding of the Fabs, cells were pelleted (7000g, 1 min) and resuspended in 4% formaldehyde (in PBS) at 4°C for further 20 minutes. Cells were then washed twice (PBS) and resuspended in a small volume of PBS. For imaging BamA migration after different growth periods, cells were grown and labelled as described above and then resuspended in fresh growth medium and grown at 37°C for the indicated time periods. OD_600_ measurements were taken immediately after resuspension in fresh medium and before each time point to validate cell growth. Wash steps were repeated as described above.

### TIRFM acquisition

Live cells were imaged using an Oxford NanoImager (ONI) superresolution microscope equipped with four laser lines (405/473/561/640 nm) and ×100 oil immersion objective (Olympus 1.49 NA). Fluorescence images were acquired by scanning a 50 μm × 80 μm area with a 473 nm laser for AF488 & GFP labelled proteins (laser power 1.4-2.3 mW) or 561 nm for mCherry labelled proteins (laser power 2.1-3.4 mW). The laser was set at 50° incidence angle (200 ms exposition), resulting in a 512 × 1024 pixel image. Images were recorded by NimOS software associated with the ONI instrument. Each image was acquired as a 20-frame stack for brightfield and fluorescence channels, respectively. For analysis, images were stacked into composite images using average intensity as a projection type in ImageJ (version 1.52p).

### FRAP experiments

FRAP was carried out using a custom-built inverted fluorescence microscope with a 100X APO TIRF, 1.45 NA oil-immersion objective lens (Nikon). A supercontinuum laser (NKT Photonics SuperK Power) provided widefield illumination with full width half maximum 14.3 μm, intensity 73 W cm^−2^ and a separately shuttered bleaching path with a width of 1 μm and intensity 10 kW cm^−2^ over a wavelength band 460-490 nm. Fluorescence images were acquired with a back-illuminated emCCD camera (Andor iXon Ultra 888) at 52 nm pixel^−1^ using a filter set comprised of band pass excitation and emission filters (469/35 and 525/39 nm), and a dichroic mirror transmitting over 498 nm. Cells were illuminated for a 100 ms exposure every 15 seconds for 10 minutes. A bleaching pulse was applied for 500 ms immediately prior to the acquisition of the second fluorescence image by camera-controlled shutters (Thorlabs SC10 and SH05).

### SIM Imaging

Cells were imaged using Deltavision OMX V3 Blaze microscopy system (GE) equipped with four laser lines (405/488/594/633 nm), a standard or a Green/Red drawer filter set and a ×60 oil immersion objective (Olympus 1.42 NA). For both conventional and SIM imaging 1.512 index refraction immersion oil was used for AF488 & GFP labelled proteins and 1.514 index refraction immersion oil was used for AF488/mCherry dual-color imaging. Conventional fluorescence images were acquired by imaging a 42 μm x 42 μm area with the 488 nm laser (5.7 mW, 500ms exposure) resulting in a 512 × 512 pixel image. For SIM acquisition, a similar area was imaged using the 488 nm laser (2.7 mW, 200 ms exposure). Image stacks of 1-1.5μm thickness were taken with 0.125 μm z-steps and 15 images (three angles and five phases per angle) per z-section and a 3D-structured illumination with stripe separation of 213 nm and 238 nm at 488 nm and 594 nm, respectively. The SIMcheck plugin (ImageJ) was used to assess the data quality of SIM images. Image stacks were reconstructed using Deltavision softWoRx 7.2.0 software with a Wiener filter of 0.003 using wavelength specific experimentally determined OTF functions. Average intensity and 3D projections of 3D-SIM images were generated using ImageJ (V1.52p)

### Single step photobleaching

For Single Step Photobleaching experiments, time lapse imaging of 300 frames (100ms each) was carried out using the Deltavision OMX V3 Blaze microscopy system (GE) with a standard drawer in conventional mode (488 nm laser, 5.7 mW). Calculating the size of a single bleaching step based on these movies was carried out as described below.

### Image and data analysis

#### Measuring the diameter of BamA islands

2D-SIM images of BamA labelled cells were binarized and Regions of interest (ROI’s) were generated. Non-distinct islands were manually excluded. The size of each island was calculated based on its Ferret’s diameter (ImageJ V1.52p).

#### Detection of islands and measuring their fluorescence intensity and distribution

Unsharp mask filter (Radius 2px, Mask weight 0.5) was applied to the raw images and BamA islands identified using the Find Maxima process and ROI’s were created. The fluorescence intensity of each ROI was measured using the raw images and background fluorescence was subtracted. For determining the distribution of BamA islands along the OM, the distance of each ROI’s from the pole was calculated and divided by the overall length of the cell. (ImageJ V1.52p).

#### Channel alignment

Alignment of dual-color images was carried out using TetraSpeck™ Microspheres, 0.1 μm (ThermoFisher scientific) and the channel aligner tool (ImageJ V1.52p).

#### Calculating the normalized fluorescence distribution of OMPs

Fluorescence distribution across the long axis of the OM was determined by the plot profile function with a line width of 3 points (ImageJ V1.52). After measuring the raw values, they were normalized to 0−1 scale for comparison between cells. To enable the integration of fluorescence intensity distribution from cells of different lengths, the long axis of each cell was normalized to 0−100. After the integration of profiles from all cells, the value at the poles was set to 1 and the rest of the values were normalized accordingly. Normalization was done using Excel and the data was plotted in GraphPad Prism 8 software.

#### Integrated localization map

For creating integrated localization maps of BamA- and LptD-containing islands, raw images were binarized to create segmented images of single cells. In parallel, islands were identified as described above and their coordinates and fluorescence intensity values were listed. Data processing and the preparation of integrated localization maps was carried out using a custom Python script. The script is available upon request from the corresponding author.

#### Calculating the size of a single bleaching step

For the analysis of BamA SSP experiments, the spot intensity analysis tool (ImageJ) was used to initially identify BamA-containing islands from which fluorescence decay curves exhibited distinct photobleaching steps. Subsequently, ROIs containing these islands were created and the fluorescence intensity trace measured. Each bleach intensity trace was then filtered using an edge-preserving Chung-Kennedy algorithm consisting of two adjacent running windows whose output was the mean from the window possessing the smallest variance (72). Photobleaching steps and blinks from multiple ROIs were used to calculate the unitary step size using a fitted Gaussian.

### Preparation of OM Extracts

50ml seed cultures were grown overnight at 37°C in LB media (unless stated otherwise) to inoculate 800ml shake flask cultures (OD_600_= 0.05). Cultures were grown with shaking at 37 °C (or the indicated temperature) until they reached mid-log phase or the required growth phase. Aztreonam was added to a final concentration of 1μg/ml 40 minutes before cells were harvested. Cell pellets were harvested by centrifugation at 5000 x g for 15 min at 4 °C and resuspended in 10 mM Tris-HCl, pH 8.0, 6 mM lithium 3,5-diiodosalicylate, 1 mM PMSF to give a final volume of 35 ml. Cells were lysed through sonication with a Misonix S4000 sonicator with insoluble cell debris being removed through centrifugation at 10,000 x g for 20 min at 4 °C. Total membrane pellets were collected from the supernatant of the low speed spin through centrifugation at 200,000g for 45 minutes at 4 °C. Inner membrane proteins were removed from the total membrane fraction through homogenising the total membrane pellet in 10 ml of 10 mM Tris-HCl, pH 8.0, 6 mM lithium 3,5-diiodosalicylate, 2 % (v/v) Triton X-100, followed by centrifugation at 200,000g for 45 minutes at 4 °C to pellet the outer membrane fraction. Outer membrane pellets were resuspended 1ml of 10 mM Tris-HCl, pH 8.0, homogenized, and analyzed on a 12 % SDS-PAGE gel.

### Western blotting

OM protein extracts were run on a 12% SDS-PAGE gel (60 mA, 30 min), volumes loaded into the gel were adjusted such that the amount of material analyzed corresponded to an equivalent OD for all samples. Subsequently, the gel was blotted onto Sequi-Blot PVDF membrane (Bio-Rad #1620182) and blocked with 8% Marvel dried skimmed milk in Tris-buffered saline buffer with Tween 20 (TBST buffer) overnight at room temperature. Blots were probed with primary rat anti-BamA (1 μg/ml) antibodies in 4% milk in TBST buffer for 1 h at room temperature. Membranes were then washed with TBST buffer (5 × 1 min) and probed with secondary rabbit anti-rat antibodies conjugated with peroxidase (1:1000, Abcam #ab6734). Blots were washed as described above and detection was carried out using Amersham ECL Western Blotting Select Detection Reagent (GE Lifesciences #RPN2235), according to the manufacturer’s instructions in GBOX-CHEMI-XRQ. Images were recorded using GeneSys software and the intensity of the bands was measured (ImageJ V1.52p).

## Acknowledgements

This work was funded by the European Research Council (Advanced grant 742555; OMPorg), the Wellcome Trust (201505/Z/16/Z) and BBSRC UK (BB/P009948/1). GM was supported by an EMBO Long-Term Fellowship (ALTF 485-2017). PGI acknowledges studentship funding from the Medical Research Council UK. We thank Emma Elliston for providing the *fepA* expressing construct and Sandip Kumar (Oxford) for help with ONI nanoimager experiments. We are also indebted to Nadia Halidi and the Micron Advanced Bioimaging Unit (Wellcome Strategic Awards 091911/B/10/Z and 107457/Z/15/Z) for their support & assistance in this work.

## Data availability

The data supporting the findings of the study are available in the article or available upon request from the corresponding author.

## Author contributions

G.M., J.P., and S.T.R. and C.K. designed research; G.M. and H.M. performed research; D.S., K.M.S., P.G.I., R.C.K., N.G.H., and R.K. contributed new reagents/analytic tools; G.M. and P.G.I. analyzed data; and G.M. and C.K. wrote the paper.

## Supplementary Figure Legends

**Fig S1.**
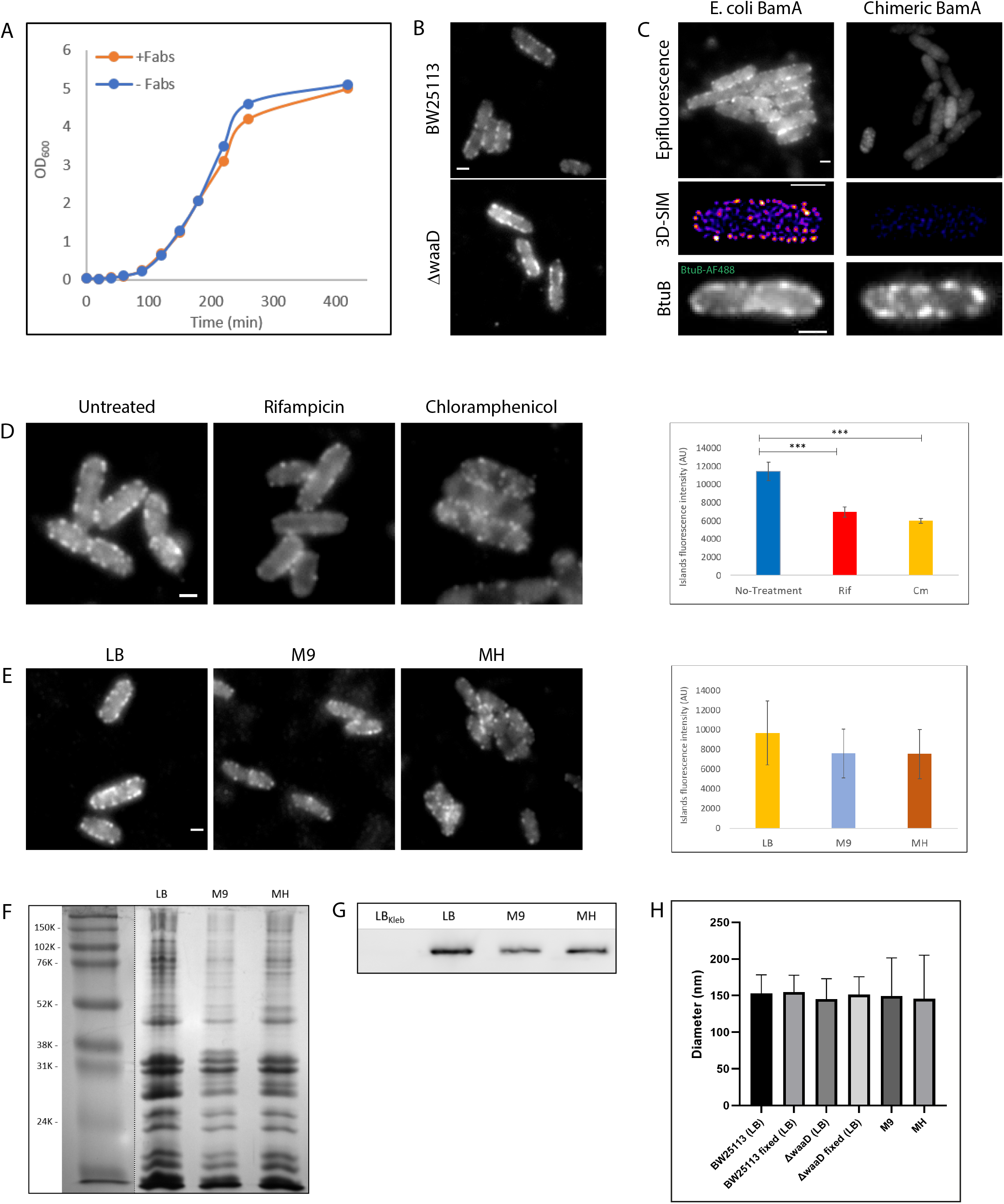
**(A)** Growth curves of *E. coli* cells grown in LB with or without Fabs used for live cell labelling. **(B)** Representative images showing a comparison of BamA labelling in the Keio wild type strain (BW25113) and the deep rough mutant strain (*waaD*). **(C)** BamA and BtuB labelling in the wild type strain and a strain expressing a modified BamA barrel domain (*K. pneumonia* sequence). Shown are comparative images of cells labelled with the indicated labels. **(D)** Epifluorescence images (*left*) and comparative BamA island fluorescence intensity (*right*) of cells treated with rifampicin and chloramphenicol. **(E)** Epifluorescence images (*left*) and comparative BamA island fluorescence intensity (*right*) of cells grown in the indicated media types. **(F)** Coomassie stained gels of OM extracts from mid-log cells grown in different media types. **(G)** BamA western blot of the OM extracts shown in panel F. The LB_Kleb_ lane was loaded with an OM extract from the strain shown in panel C (negative control). **(H)** The average diameter of BamA-containing islands in different strains, growth media and after fixation. Measurements are based on 2D-SIM images. Scale bars, 1μm.

**Fig S2.**
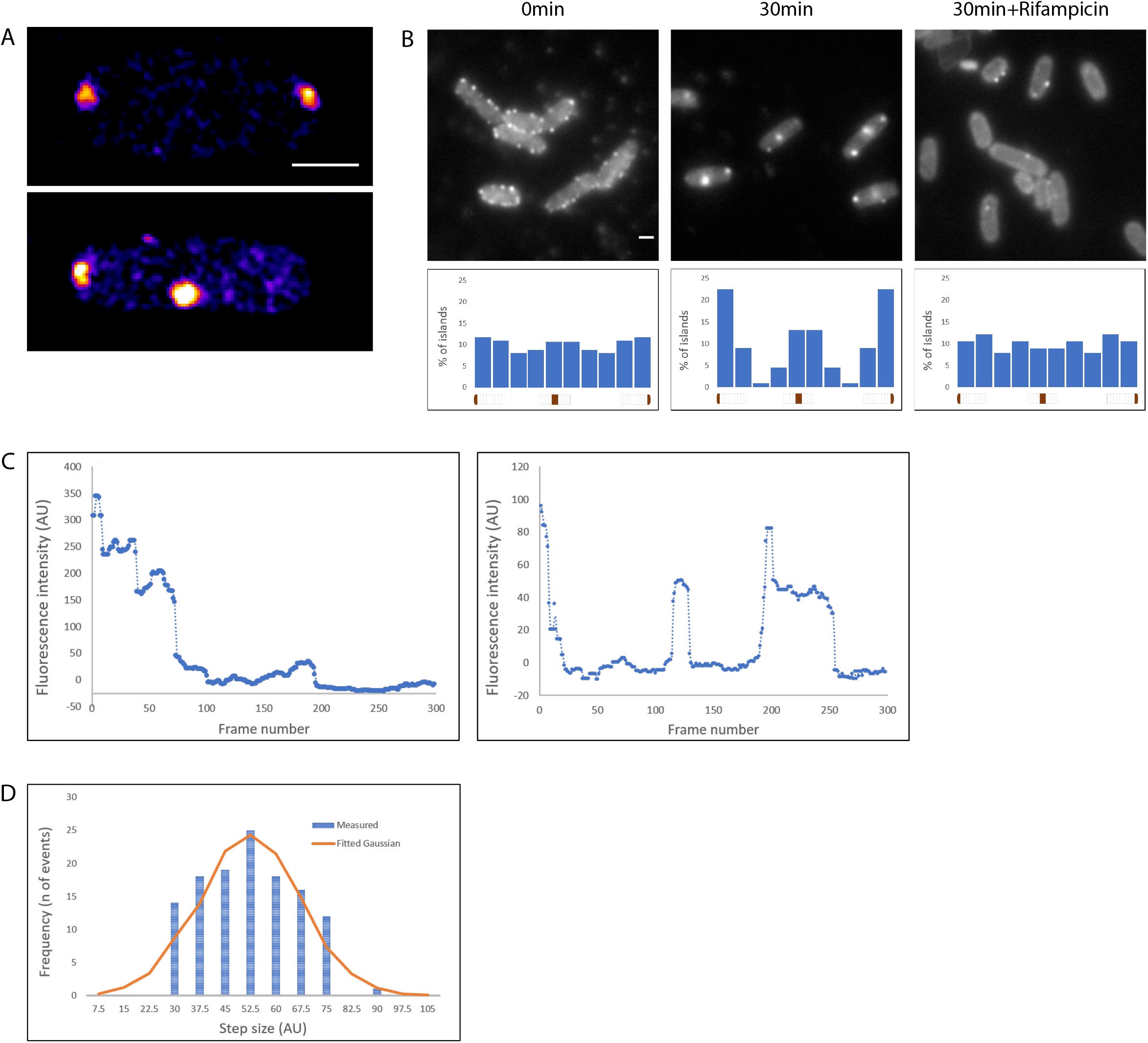
**(A)**3D-SIM projections (heatmaps) of BamA migration to the poles after 30 minutes of growth. **(B)***E. coli* cells were labelled with αBamA^−AF488^ Fabs at T_0_, and resuspended in fresh LB media +/− rifampicin for 30 minutes. Shown are representative images of BamA-containing islands (*top panels*) and their distribution plots (*bottom panels*). (n_T0_=132, n_T30_=111, n_T30rif_=95). **(C)** Representative fluorescence decay curves from Single Step Photobleaching experiments. Shown are examples of stepped signal decay (*left*) and blinks of single molecules (*right*) after filtering the raw data with the Chung–Kennedy algorithm. **(D)** Distribution of step sizes from multiple SSP experiments (n=123). The size of a single photobleaching step was calculated based on a fitted Gaussian. Scale bars, 1 μm.

**Fig S3.**
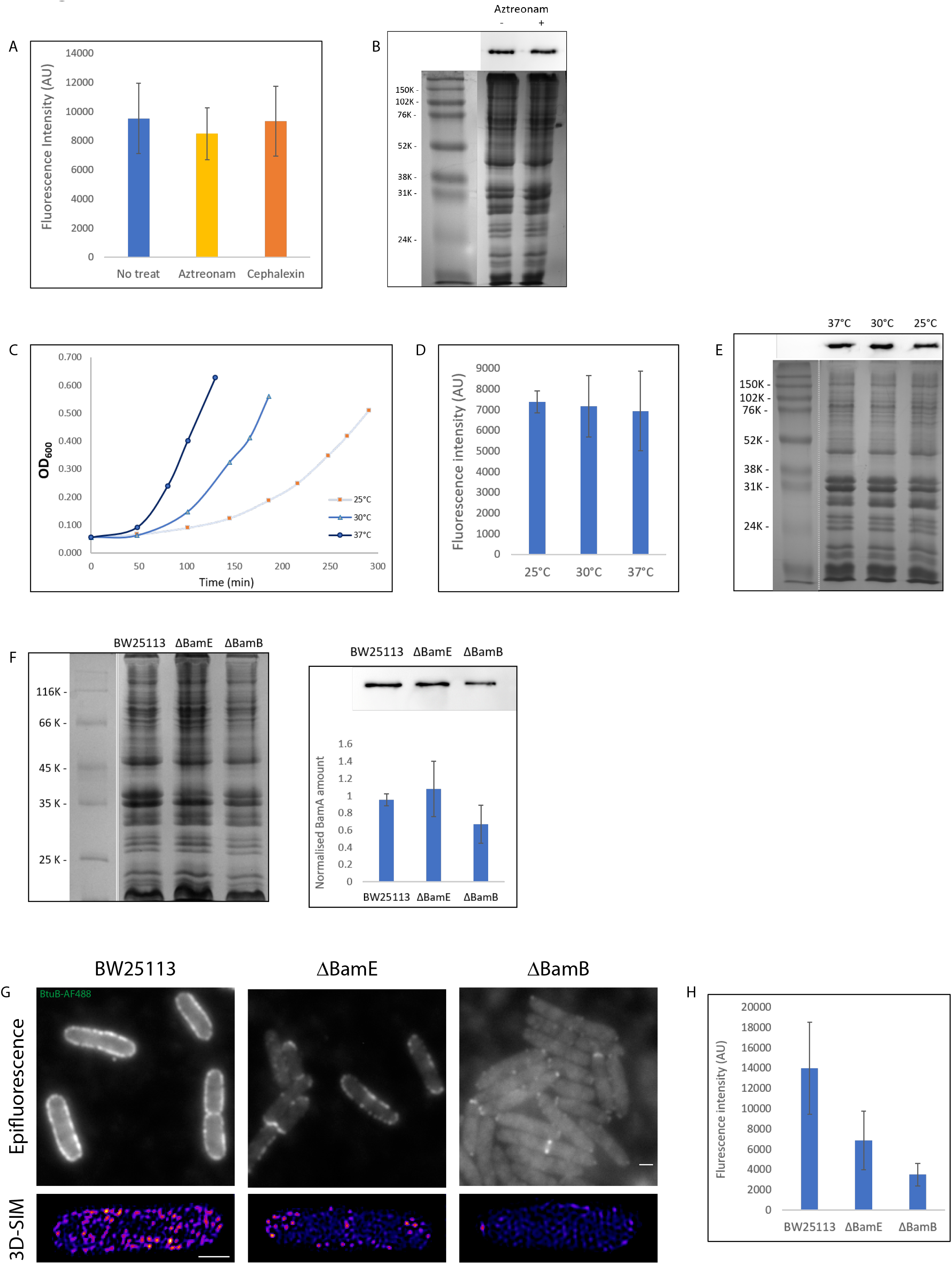
**(A)** Fluorescence intensity of BamA-containing islands in cells treated with antibiotics inhibiting cell division. **(B)** Coomassie blue-stained SDS gel and BamA Western blot of OM extracts from cells grown in LB media with or without aztreonam treatment. **(C)** Growth curves of *E. coli* cultures grown in LB at different temperatures. For each culture, samples were taken for labelling and imaging (Figure 2C) at the last time point. **(D)** Fluorescence intensity of BamA-containing islands in cells grown at different temperatures. **(E)** Coomassie blue-stained SDS gel and BamA western blot of OM extracts from cells grown at different temperatures. **(F)** Coomassie stained SDS gel of OM extracts from wild type (BW25113), *bamE* and *bamB* strains. **(G)** BamA western blot of OM extracts from the indicated strains. Shown is a representative blot and the quantification from four biological replicates. **(H)** Epifluorescence and 3D-SIM images (heatmaps) of BtuB labelling in the indicated strains. **(I)** Fluorescence intensity of BtuB within islands in the indicated strains. Scale bars, 1μm.

**Fig S4.**
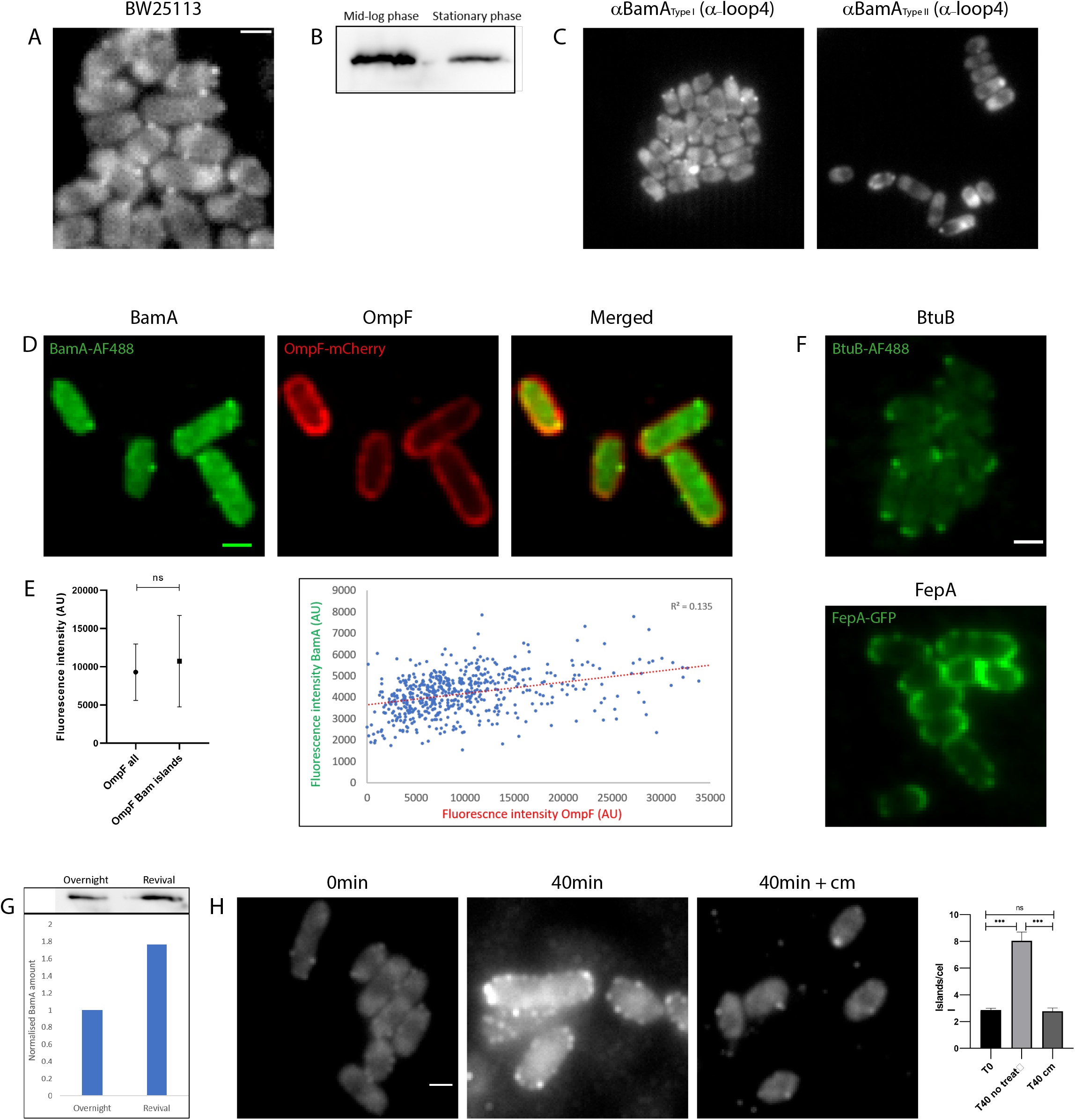
**(A)** Representative epifluorescence image of BamA labelled at stationary phase (BW25113). **(B)** BamA western blot of OM extracts from mid-log vs stationary phase cells. Sample loading was based on equivalent OD_600_ measurements. **(C)** Images of BamA labelling at stationary phase using MAB1 αBamA Fab, which binds loop 4 (15). **(D)** Co-labelling of BamA and OmpF in stationary phase cells. **(E)** Comparison of OmpF fluorescence intensity in the entire cell vs regions containing BamA-containing islands (*left*) and a scatter plot showing the lack of correlation between fluorescence intensity of BamA and OmpF (*right*). **(F)** Representative fluorescence images of BtuB and FepA in stationary phase cells (BW25113). **(G)** BamA western blot of OM extracts from an overnight culture vs revived cells (cells were resuspended in fresh LB until the OD_600_ raised from 0.050 to 0.075). Sample loading was OD_600_ adjusted. **(H)** Fluorescence images of labelled BamA after resuspension in fresh media with or without chloramphenicol treatment. Shown are representative images (*left*) and the number of islands per cell detected by epifluorescence microscopy (*right*). Scale bars, 1 μm

**Fig S5.**
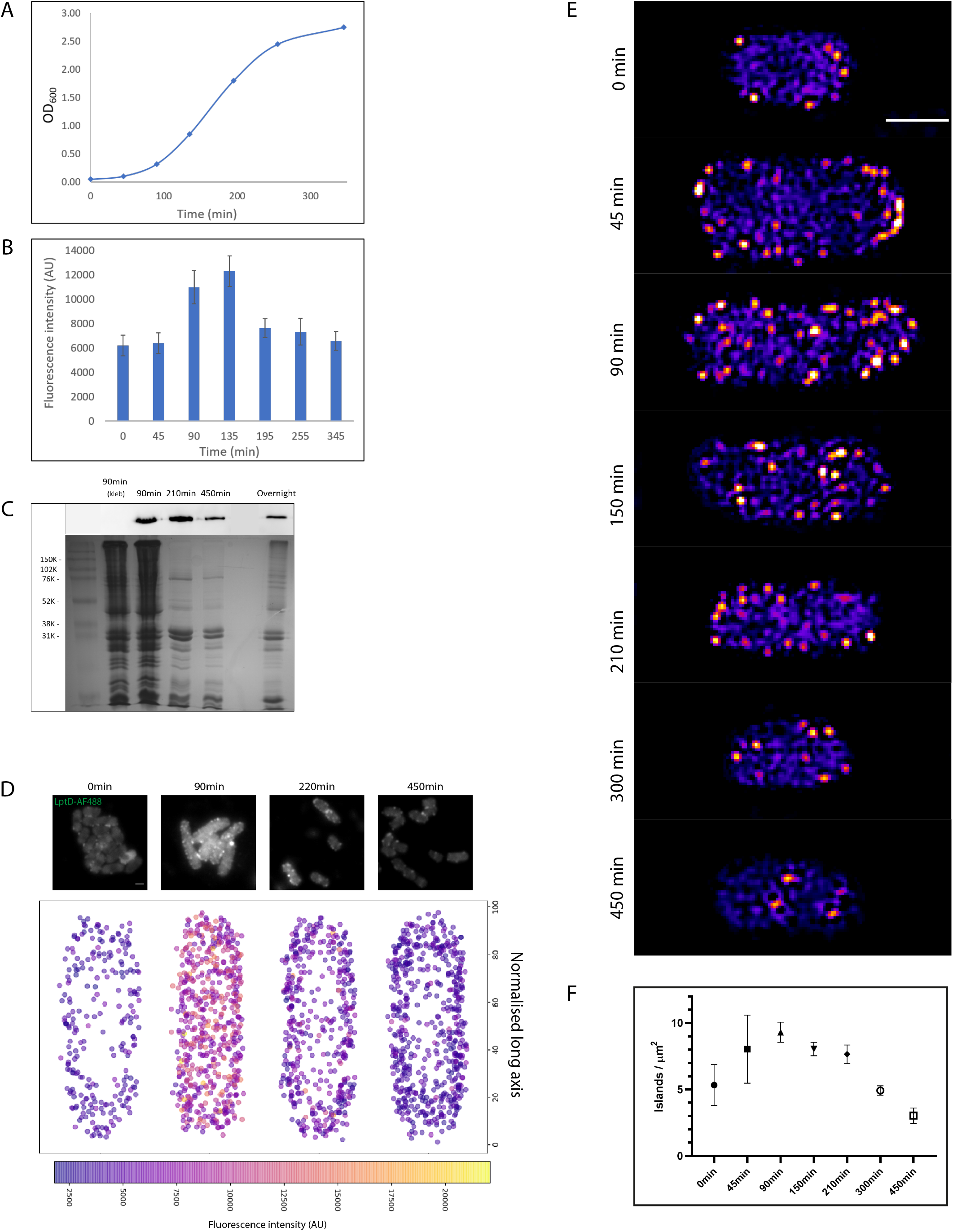
**(A)** Growth curve of *E. coli* cells grown for the BamA time course experiment (Figure 4F). **(B)** Fluorescence intensity of BamA islands at different growth stages. **(C)** Coomassie stained SDS gel and a BamA western blot of OM extracts from cells at different growth stages (90min_kleb_ is a negative control). **(D)** Time-course images and integrated localization maps of LptD islands at different growth stages. Islands are color coded according to their fluorescence intensity. Scale bars, 1 μm **(E)** Representative 3D-SIM images of BamA labelling at the indicated time points analyzed in panel D. **(F)** Density of BamA islands at 7 different time points during a time-course experiment. Quantifications are based on 3D-SIM images. Time-course images and integrated localization maps of LptD islands at different growth stages. Islands are color coded according to their fluorescence intensity. Scale bars, 1 μm

**Fig S6.**
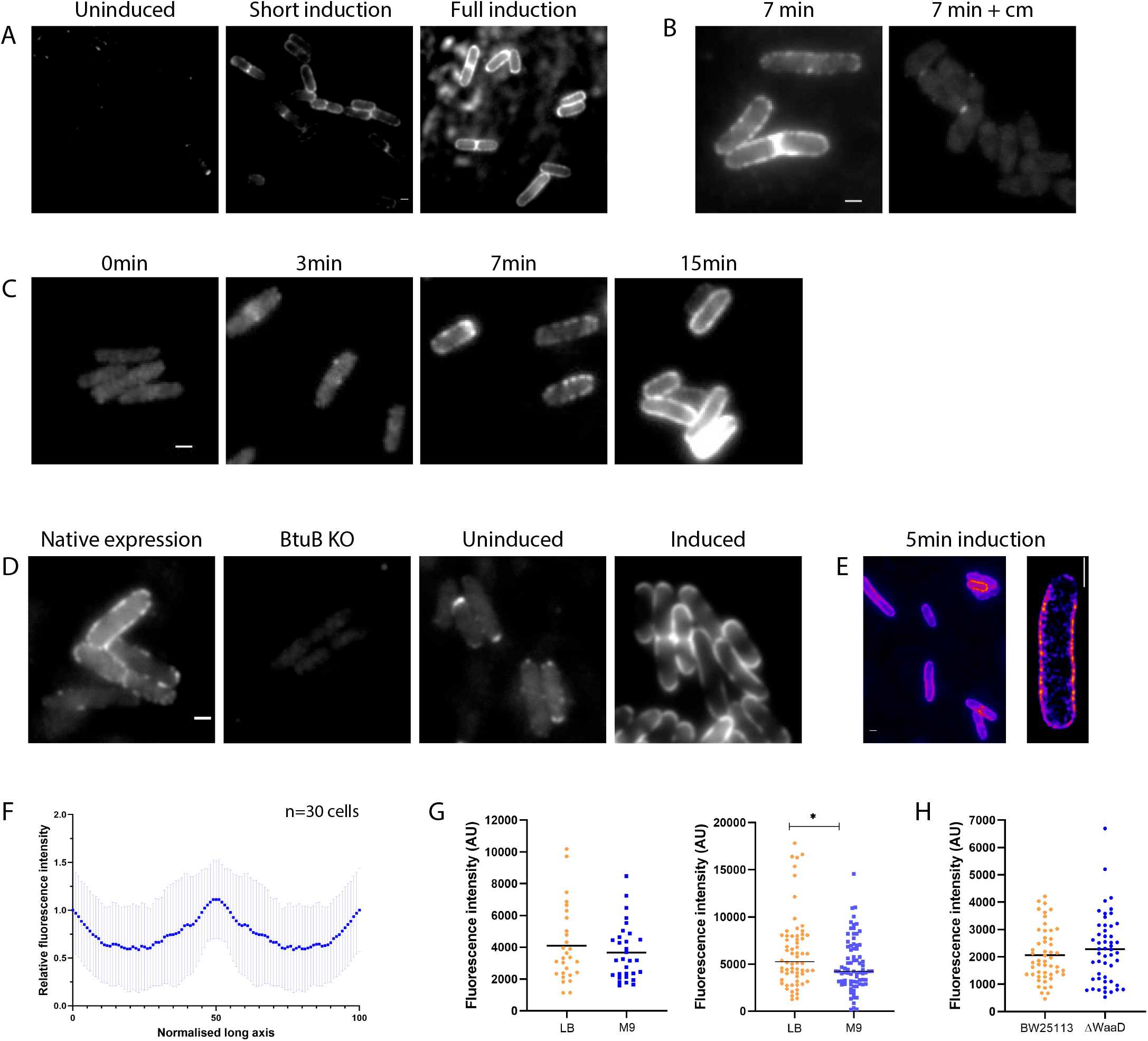
**(A)** Epifluorescence images of FepA in an inducible expression strain. The images shown are of labelled cells after different induction regimes. **(B)** Images of FepA labelling after 7 minutes induction with or without chloramphenicol pre-treatment. **(C)** Timeline of FepA biogenesis in M9 media. Samples were taken and labelled at several time points after FepA induction. **(D)** Epifluorescence images of BtuB in the indicated strains and under different induction regimes. (**E)** Representative images of mid-cell biased BtuB biogenesis after 5 minutes of BtuB induction. Shown are epifluorescence and 3D-SIM projection heatmaps. **(F)** Kymograph of normalized fluorescence intensity profiles based on the integrated analysis of multiple cells 5 minutes after BtuB induction. **(G)** Comparison of FepA (*left*) and BtuB (*right*) fluorescence intensity after 5 minutes induction in LB vs. M9. **(H)** Comparison of FepA fluorescence intensity after 5 minutes induction in wild type (BW25113) cells vs the LPS deep rough mutant strain (*waaD*). Scale bars, 1 μm.

**Fig S7.**
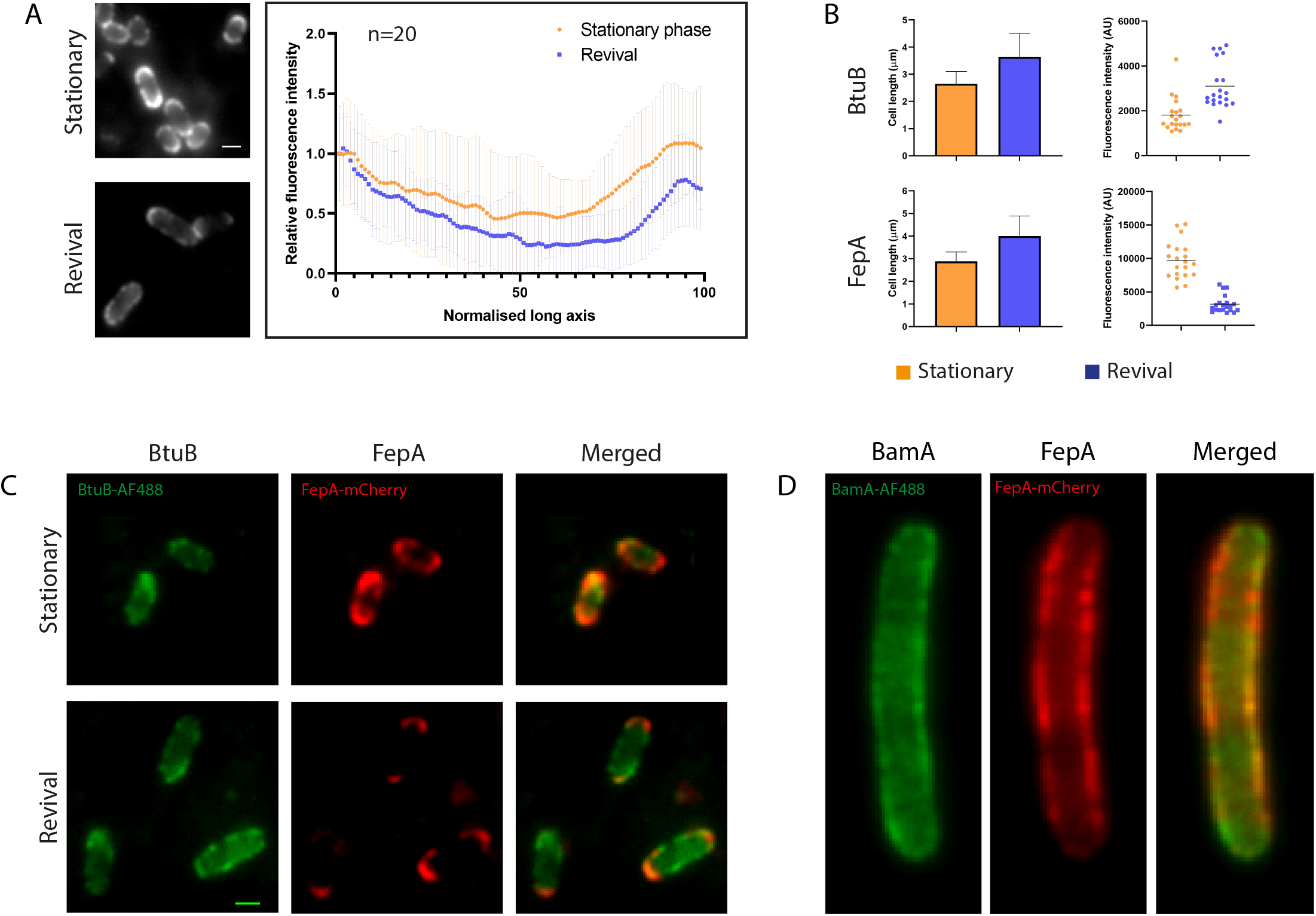
**(A)** Representative images (*left*) and a kymograph (*right*) of FepA fluorescence intensity profiles at stationary phase vs revival in fresh media. **(B)** Charts comparing the average length and fluorescence intensity of BtuB and FepA in stationary phase before vs after a short revival in fresh media. **(C)** Co labelling of BtuB and FepA in stationary phase and after a short revival in fresh media. **(D)** Co-labelling of BamA and FepA after 7 minutes induction of FepA. Shown are epifluorescence images of a long cell. Scale bars, 1 μm

**Table S1.**
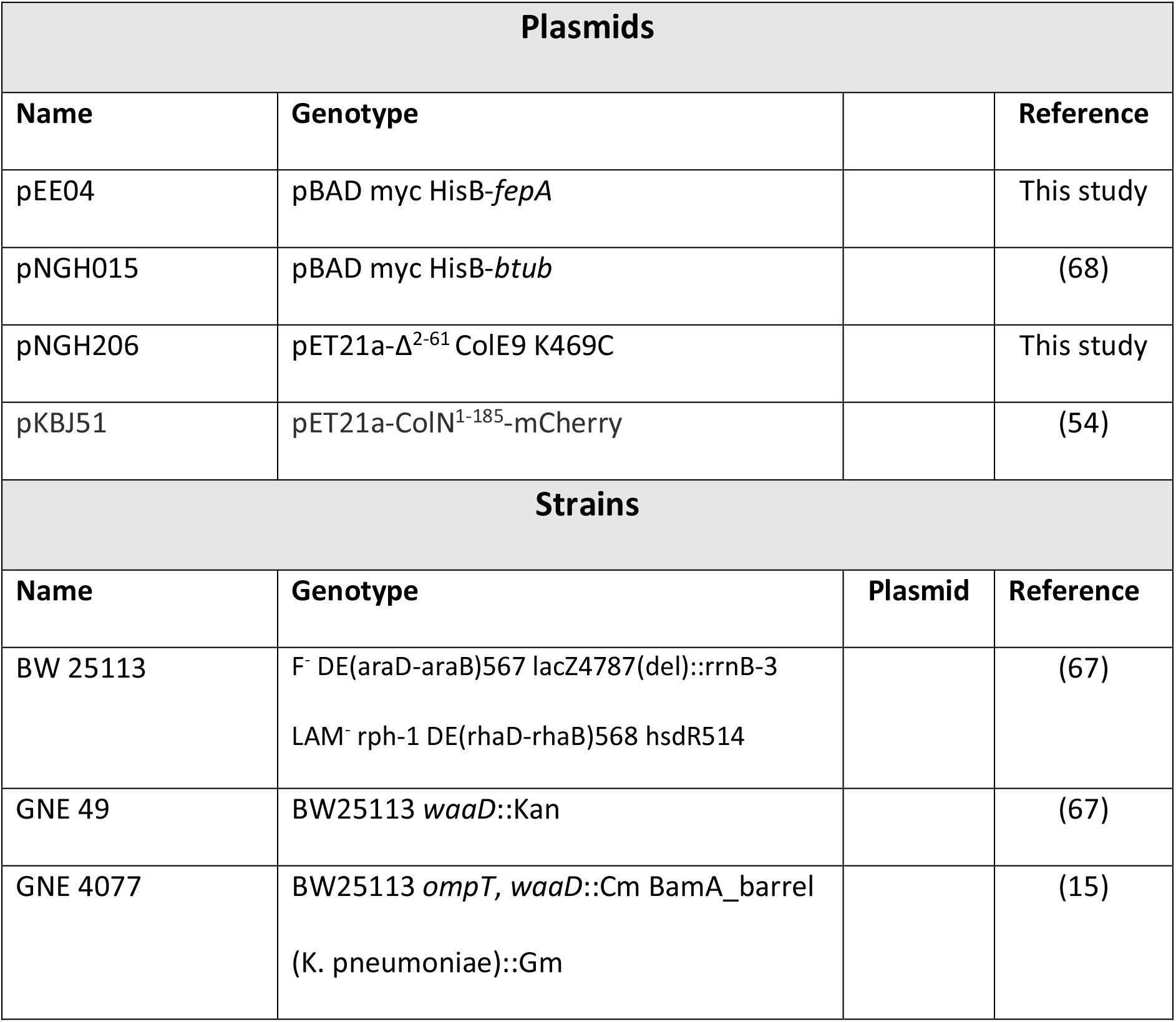
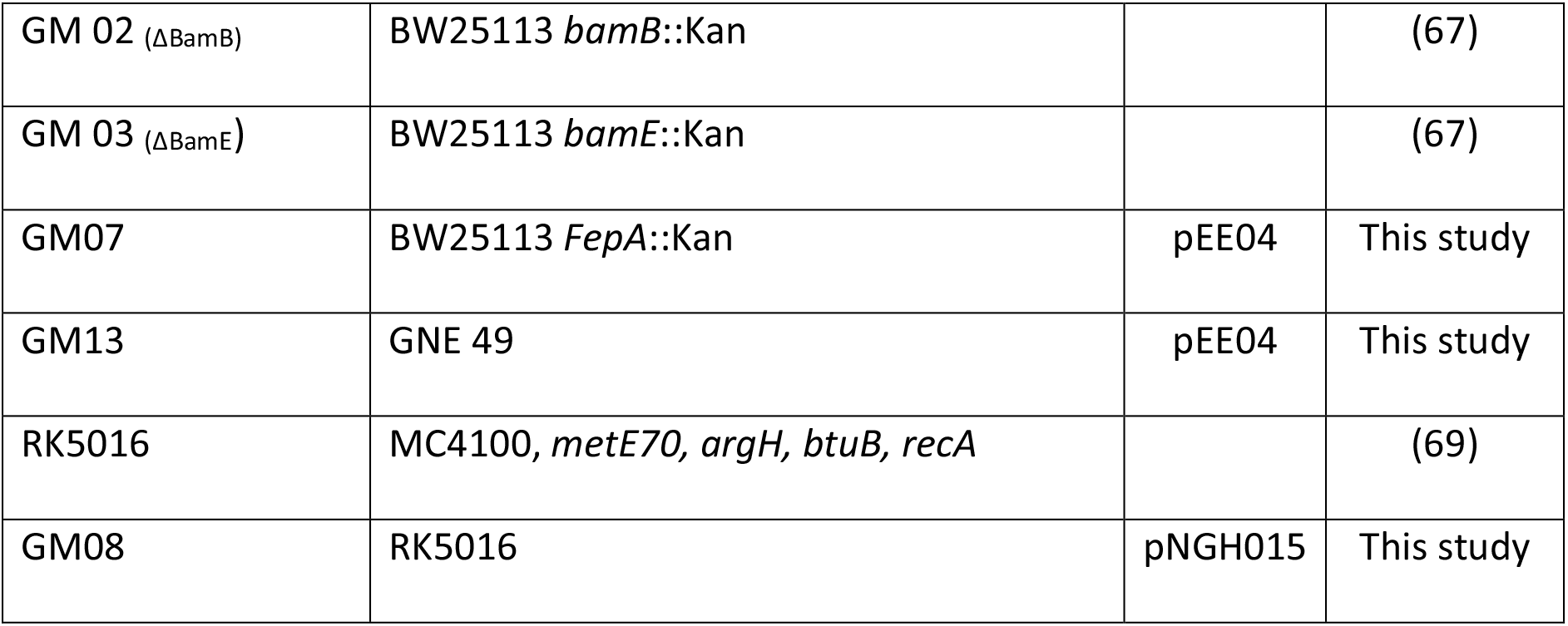

| Plasmids |  |  |  |
| --- | --- | --- | --- |
| Name | Genotype |  | Reference |
| pEE04 | pBAD myc HisB-*fepA* |  | This study |
| pNGH015 | pBAD myc HisB-*btub* |  | (68) |
| pNGH206 | pET21a-$\Delta^{2-61}$ ColE9 K469C | | This study |
| pKBJ51 | pET21a-ColN<sup>1-185</sup>-mCherry |  | (54) |
| Strains |  |  |  |
| Name | Genotype | Plasmid | Reference |
| BW 25113 | F<sup>-</sup> DE(araD-araB)567 lacZ4787(del)::rrnB-3<br>LAM<sup>-</sup> rph-1 DE(rhaD-rhaB)568 hsdR514 |  | (67) |
| GNE 49 | BW25113 *waaD*::Kan |  | (67) |
| GNE 4077 | BW25113 *ompT*, *waaD*::Cm BamA_barrel<br>(*K. pneumoniae*)::Gm |  | (15) |
| GM 02 (ΔBamB) | BW25113 *bamB*::Kan | | (67) |
| GM 03 (ΔBamE) | BW25113 *bamE*::Kan | | (67) |
| GM07 | BW25113 *FepA*::Kan | pEE04 | This study |
| GM13 | GNE 49 | pEE04 | This study |
| RK5016 | MC4100, *metE70*, *argH*, *btuB*, *recA* |  | (69) |
| GM08 | RK5016 | pNGH015 | This study |

**Table S2.**

| Name | Type | Reference |
| --- | --- | --- |
| αBamA Abs (western blots) | Ab, monoclonal | (15) |
| αBamA 29e9 (MAB2) | Fab, monoclonal | (15) |
| αBamA 15c9 (MAB1) | Fab, monoclonal | (15) |
| αLptD | Fab, monoclonal | (71) |
| ColE9-AF488 | Colicin | (44) |
| ColB-GFP/mCherry | Colicin | This study |
| ColN-mCherry | Colicin | (54) |

## Movies legends

**Movie S1** - 3D image of BamA labelling using monoclonal Fabs (heatmap) and the binary map of the islands.

**Movie S2** - BamA FRAP movie.

**Movie S3** - BamA FRAP movie, bleaching of a polar position.

**Movie S4** – A representative SSP movie used to identify photobleaching steps and blinks.

**Movie S5** - 3D image of BamA labelling (heatmap) in the *bamB* and *bamE* strains.

**Movie S6** - 3D image of LptD labelling using monoclonal Fabs (heatmap)

**Movie S7** - 3D image of FepA labelling after 5 minutes induction (heatmap).

**Movie S8** - 3D image of BtuB after 5 minutes induction (heatmap).

**Movie S9** - 3D images of BtuB at stationary phase and after revival.

**Movie S10** - 3D images of FepA at stationary phase and after revival.

